# Mechanistic and evolutionary insights into isoform-specific ‘supercharging’ in DCLK family kinases

**DOI:** 10.1101/2023.03.29.534689

**Authors:** Aarya Venkat, Grace Watterson, Dominic P. Byrne, Brady O’Boyle, Safal Shrestha, Nathan Gravel, Emma E. Fairweather, Leonard A. Daly, Claire Bunn, Wayland Yeung, Ishan Aggarwal, Samiksha Katiyar, Claire E. Eyers, Patrick A. Eyers, Natarajan Kannan

## Abstract

Catalytic signaling outputs of protein kinases are dynamically regulated by an array of structural mechanisms, including allosteric interactions mediated by intrinsically disordered segments flanking the conserved catalytic domain. The Doublecortin Like Kinases (DCLKs) are a family of microtubule-associated proteins characterized by a flexible C-terminal autoregulatory ‘tail’ segment that varies in length across the various human DCLK isoforms. However, the mechanism whereby these isoform-specific variations contribute to unique modes of autoregulation is not well understood. Here, we employ a combination of statistical sequence analysis, molecular dynamics simulations and *in vitro* mutational analysis to define hallmarks of DCLK family evolutionary divergence, including analysis of splice variants within the DCLK1 sub-family, which arise through alternative codon usage and serve to ‘supercharge’ the inhibitory potential of the DCLK1 C-tail. We identify co-conserved motifs that readily distinguish DCLKs from all other Calcium Calmodulin Kinases (CAMKs), and a ‘Swiss-army’ assembly of distinct motifs that tether the C-terminal tail to conserved ATP and substrate-binding regions of the catalytic domain to generate a scaffold for auto-regulation through C-tail dynamics. Consistently, deletions and mutations that alter C-terminal tail length or interfere with co-conserved interactions within the catalytic domain alter intrinsic protein stability, nucleotide/inhibitor-binding, and catalytic activity, suggesting isoform-specific regulation of activity through alternative splicing. Our studies provide a detailed framework for investigating kinome–wide regulation of catalytic output through cis-regulatory events mediated by intrinsically disordered segments, opening new avenues for the design of mechanistically-divergent DCLK1 modulators, stabilizers or degraders.

## Introduction

Protein kinases are one of the largest druggable protein families comprising 1.7% of the human genome and play essential roles in regulating diverse eukaryotic cell signaling pathways (1). The Doublecortin-like kinases (DCLKs) are understudied members of the calcium/calmodulin-dependent kinase (CAMK) clade of serine-threonine kinases (2–4). There are three distinct paralogs of DCLK (1, 2, and 3), the last of which is annotated as a “dark” kinase due to the lack of information pertaining to its function (5). Human DCLK1 (also known as DCAMKL1) was initially identified in 1999 (6), followed by the cloning of human DCLK2 and 3 paralogs (7). Full-length DCLK proteins contain N-terminal Doublecortin-like (DCX) domains, microtubule-binding elements that play a role in microtubule dynamics, neurogenesis, and neuronal migration (8,9). DCLKs have garnered much interest as disease biomarkers, since they are upregulated in a variety of cancer pathologies (10–12), as well as neurodegenerative disorders such as Huntington’s Disease (13). However, the mechanisms by which DCLK activity is auto-regulated, and how and why they have diverged from other protein kinases is not well understood.

Like all protein kinases, the catalytic domain of DCLKs adopts a bi-lobal fold (4), with an N-terminal ATP binding lobe and C-terminal substrate binding region. Canonical elements within the two lobes include the DFG motif, a Lys-Glu salt bridge that is associated with the active conformation, Gly-rich loop, and ATP-binding pocket, which are all critical elements for catalysis. Many protein kinases, including CAMKs, Tyrosine Kinases (TKs) and AGCs, elaborate on these core elements with unique N-terminal and C-terminal extensions that flank these catalytic lobes (14–18), allowing them to function as allosteric regulators of catalytic activity (4). Indeed, CAMKs are archetypal examples of kinases that can exist in an active-like structural conformation, yet still remain catalytically inactive (4). This is in large part due to the presence of unique C-terminal tails that are capable of blocking ATP or substrate binding in well-studied kinases such as CAMK1 and CAMKII. In canonical CAMKs, autoinhibition may be released upon Ca^2+^/Calmodulin (CaM) interaction with the CAMK C-tail, which makes the substrate-binding pocket and enzyme active-site accessible (19). In CAMKII, the N and C-terminal segments flanking the kinase domain are variable in length across different isoforms and the level of kinase autoinhibition or autoactivation has been reported to be dependent on the linker length (20). The CAMKII C-tail can be organized into an autoregulatory domain and an intrinsically disordered association domain. The autoregulatory domain also serves as a pseudosubstrate, which physically blocks the substrate binding pocket until it is competed away by CaM (21). Notably, this autoregulatory pseudosubstrate can be phosphorylated (19), and phosphorylation of the C-tail makes CAMKII insensitive to CaM binding. Across the CAMK group, several other kinases share autoinhibitory activity via interactions between Ca^2+^/CaM binding domains and the C-terminal tail (22,23), and a major feature of these kinases is variation in the tail length across the distinct genetic isoforms.

The human genome encodes four distinct DCLK1 isoforms, termed DCLK1.1-1.4 in UniProt (**Table 1**, **Figure 1**, (24)), which display differential activity– and tissue-specific expression profiles. Human DCLK1.1 (also known as DCLK1 alpha) is expressed in a variety of tissues, but is enriched in cells derived from the fetal and adult brain, whereas DCLK1.2 (also known as DCLK1 beta) is expressed exclusively in the embryonic brain (25). DCLK1.3 and 1.4, which lack tandem microtubule-binding DCX domains (**Figure 1**) but are otherwise identical to DCLK1.1 and DCLK1.2, respectively, are also highly expressed in the brain. To aid with clarity, the names of the human DCLK1 genes and their isoforms used in this paper are summarized in **Table 1**. Recent structural and cellular analyses have begun to clarify the mechanisms by which the DCLK1.2 isoform is regulated by the C-tail (2,26,27). Mechanistically, autophosphorylation of Thr 688, which is present only in the C-tail of DCLK1.2 (and DCLK1.4), blunts kinase activity and subsequently inhibits phosphorylation of the N-terminal DCX domain and thus drives DCLK microtubule association in cells (2). Consistently, deletion of the C-tail or mutation of Thr 688 restores DCLK1.2 kinase activity, subsequently leading to DCX domain phosphorylation and the abolition of microtubule binding. The length and sequence of the C-tail varies across the DCLK1 isoforms; however, how these variations contribute to isoform-specific functions and how they emerged during the course of evolution is not known.

**Figure 1:**
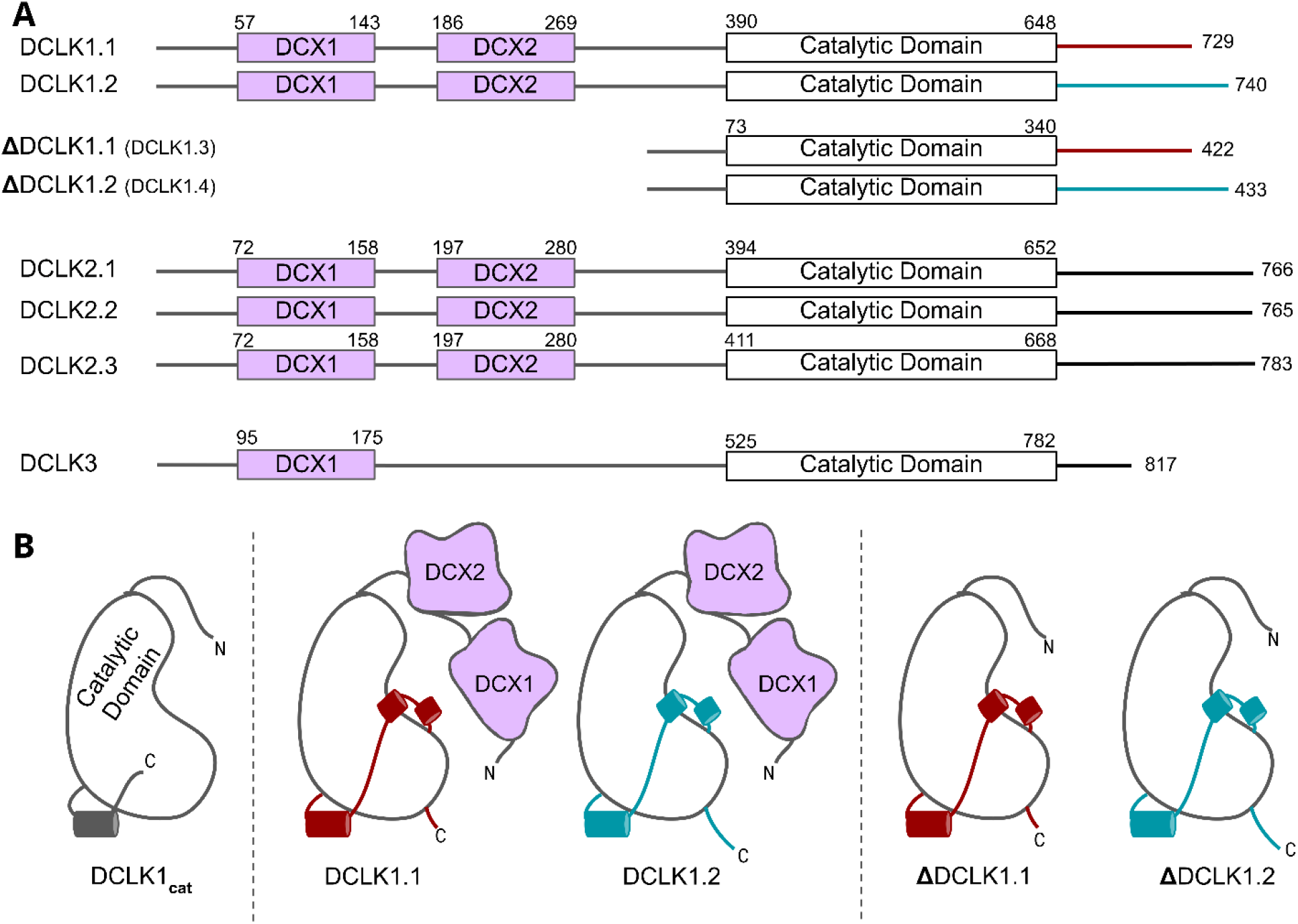
A) Schematic representation of domain organization for the known isoforms of the three human DCLK paralogs. Domain boundaries are annotated according to the representative amino acid sequences derived from UniProt. **B)** DCLK1 isoforms visualized as cartoons, showing key structural differences between the four human DCLK1 isoforms and a DCLK1 catalytic domain with artificially short linker regions (DCLK1_cat_).

**Table 1:** Names of DCLK isoforms discussed in this paper, along with their respective isoform number, UniProt identification, and alternate names that have been used.

| Name (This Study) | Isoform number | UniProt ID | Alternate names |
| --- | --- | --- | --- |
| DCLK1.1 | 1 | O15075-2 | DCAMKL1 alpha |
| DCLK1.2 | 2 | O15075-1 | DCAMKL1 beta |
| ΔDCLK1.1 | 3 | O15075-3 |  |
| ΔDCLK1.2 | 4 | O15075-4 |  |

In this paper, we employ an evolutionary systems approach that combines statistical sequence analysis with experimental studies to generate new models of DCLK evolutionary divergence and functional specialization. We identify the C-terminal tail as the hallmark of DCLK functional specialization across the kingdoms of life and propose a refined model in which this regulated tail functions as a highly adaptable ‘Swiss-Army knife’ that can ‘supercharge’ multiple aspects of DCLK signaling output. Notably, a conserved segment of the C-tail functions as an isoform-specific autoinhibitory motif, which mimics ATP functions through direct tail docking to the nucleotide-binding pocket, where it forms an ordered set of interactions that aligns the catalytic (C) spine of the kinase in the absence of ATP binding. Furthermore, molecular modelling demonstrates that a phosphorylated threonine in the C-tail of DCLK1.2, which is absent in DCLK1.1, is positionally-poised to competitively mimic the gamma phosphate of ATP, perhaps in a regulated manner. Other segments of the tail function as a pseudosubstrate by occluding the substrate-binding pocket and tethering to key functional regions of the catalytic domain. Thermostability analysis of purified DCLK1 proteins, combined with molecular dynamics simulations, confirms major differences in thermal and dynamic profiles of the DCLK1 isoforms, while catalytic activity assays reveal how specific variations in the G-loop and C-tail can rescue DCLK1.2 from the autoinhibited conformation. Together, these studies demonstrate that isoform-specific variations in the C-terminal tail co-evolved with residues in the DCLK kinase domain, contributing to regulatory diversification and functional specialization.

## Results

### Origin and evolutionary divergence of DCLK family members

The human DCLKs repertoire is composed of three genes, termed DCLK1, 2 and 3 (**Figure 1A, Table 1**). The experimental model employed in this study, DCLK1, is composed of multiple spliced variants in human cells. Those full-length proteins that contain N-terminal DCX domains are usually referred to as DCLK1.1 or DCLK1.2 and the variants that lack the DCX domains are termed here (for simplicity) ΔDCLK 1.1 and ΔDCLK1.2 (also referred to as DCLK1.3 and DCLK1.4). The core catalytic domain with minimal flanking regions (DCLK1_cat_, **Figure 1B**) is identical in all DCLK1 proteins, whereas the length of the tail, or the presence of the DCX domains, generates considerable diversity from the single human DCLK1 gene (**Figure 1B, Figure 1-figure supplement 1**). To infer evolutionary relationships of DCLK paralogs, and especially the evolution of the C-terminal tail regions that lie adjacent to the kinase domain (**Figure 1**), we performed phylogenetic analysis of 36 DCLK sequences with an outgroup of closely related CAMK sequences (**Figure 2A, Figure 2-source data 1**). These DCLK sequences are from a representative group of holozoans, which consist of multicellular eukaryotes (metazoans) and closely related unicellular eukaryotes (pre-metazoans). The analysis generated four distinct clades: pre-metazoan DCLK, metazoan DCLK3, vertebrate DCLK2 and vertebrate DCLK1. Interestingly, DCLK genes demonstrated significant expansion and diversification within metazoan taxa. The pre-metazoan DCLK sequences were the most ancestral and showed no DCLK diversity, suggesting the DCLK expansion and diversification correlated with the evolution of multicellular organisms. Within the metazoan expansion of DCLK, DCLK3 is the most ancestral and can be broken down into two sub-clades: protostome DCLK3 and deuterostome DCLK3. Within invertebrates, only two DCLK paralogs were present, one that was identified as a DCLK3 ortholog and another that was not clearly defined as either DCLK1 or DCLK2. This suggests that the diversification into DCLK1 and DCLK2 paralogs from an ancestral DCLK1/2-like paralog occurred after the divergence of invertebrates and vertebrates, which is further supported by the monophyletic DCLK1 and DCLK2 clades in vertebrates (bootstrap value: 99).

**Figure 2:**
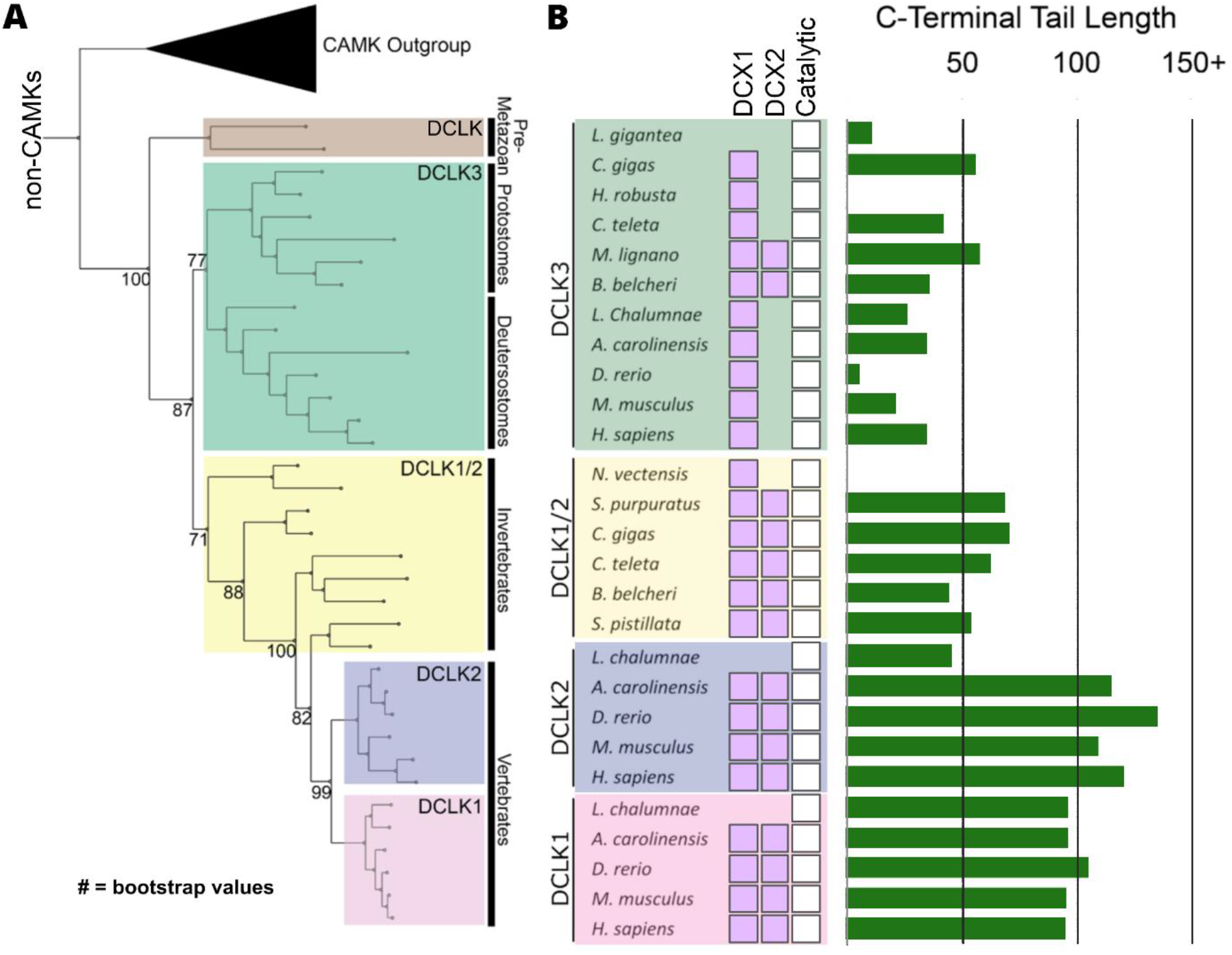
Evolution of the DCLK family. A) Phylogenetic tree showing the divergence and grouping of DCLK sub-families in different taxonomic groups. Bootstrap values are provided for each clade. **B)** Shows domain annotations for sequences included in the phylogenetic tree. The length of C-terminal tail segment for these sequences is shown as a histogram (green). The original tree generated using IQTREE is provided in Figure 2**-source data 1**.

Interestingly, the expansion of DCLK in metazoans and the diversification of DCLK1 and DCLK2 within vertebrates correlates well with the length and sequence similarity of the C-terminal tail, which also varies between the different DCLK1 splice variants (**Figure 1A, Figure 1-figure supplement 1 and 2**). Within both protostome and deuterostome DCLK3, the length of the C-terminal tail is ∼50 residues or less. This is in marked contrast to the tail lengths of vertebrate DCLK1 and DCLK2, which are ∼100 residues long. In addition to the C-terminal tail, an analysis of the domain organization of these DCLKs reveals that DCLK3 predominantly contains only a single N-terminal Doublecortin domain (DCX), whereas invertebrate DCLKs, and vertebrate DCLK1 and DCLK2 predominantly contain two DCX domains at the N-terminus of the long isoforms (**Figure 2B**). In addition, we identified a putative active site-binding motif, VSVI, and a phosphorylatable threonine conserved within vertebrate DCLK1 and DCLK2, which is absent in all other DCLK sequences, including invertebrate DCLK1/2. This raises the possibility that the DCLK1/2 tail extensions are employed for vertebrate-specific regulatory functions.

Next, we compared the type of DCLK1 protein sequence encoded by a range of chordate mammalian genomes. The domain organization of each DCLK1 isoform was compared based on annotated sequences from UniProt, demonstrating the presence of at least one DCLK1 protein that lacks the DCX domains in every species examined, with a mixture of ΔDCLK1.1 and ΔDCLK1.2 splice variants. Interestingly, it was only in the human DCLK1 gene that definitive evidence for ΔDCLK1.1 and ΔDCLK1.2 variants erewere found (**Figure 3A**). To establish a model for DCLK1 biophysical analysis, we constructed a recombinant hybrid human DCLK1 catalytic domain with a short C-tail sequence that is equivalent to DCLK1.1 amino acids 351-689, containing the catalytic domain with a short C-tail region. As shown in **Figure 3B**, incubation of size-exclusion chromatography (SEC) purified GST-tagged DCLK1 with 3C protease generated the mature untagged DCLK1 protein for biophysical analysis. Analytical SEC revealed that purified DCLK1.1 and DCLK1.2 isoforms are monomeric in solution (**Figure 3-figure supplements 1-3**). We evaluated catalytic activity for DCLK1.1_351-689_ using a validated peptide phosphorylation assay (**Figure 3C**), which revealed efficient phosphorylation of a DCLK1 substrate peptide. The *K*_M[ATP]_ for peptide phosphorylation was close to 20 µM in the presence of Mg^2+^ ions (**Figure 3C**, left panel), similar to values measured for other Ser/Thr kinases that are autophosphorylated and active after expression from bacteria (28). DCLK-dependent peptide phosphorylation was completely blocked (**Figure 3C**, right panel) by prior incubation of the reaction mixture (containing 1mM ATP) with the chemical inhibitor DCLK1-IN-1, as expected (29). In addition to enzyme activity, we monitored thermal denaturation of purified, folded, DCLK1_351-689_ protein in the presence of ATP, either alone or as a Mg:ATP complex, which is required for catalysis. As shown in **Figure 3D**, DCLK1 was stabilized by 2.1°C upon incubation with an excess of Mg:ATP, and this protective effect was completely blocked by mutation of Asp 533 (of the conserved DFG motif) to Ala, consistent with canonical ATP interaction in the nucleotide-binding site. Finally, we assessed the thermal effects of a panel of DCLK1 inhibitors on the model DCLK1.1_351-689_ protein. Prior incubation with DCLK1-IN-1, LRRK2-IN-1, the benzopyrimidodiazipinones XMD8-92 and XMD8-85, which have been reported to potently (though not specifically) inhibit DCLK1 activity (26), led to marked protection from thermal unfolding (**Figure 3-figure supplement 4**). Consistently, the negative control compound DCLK1-Neg (29) was ineffective in stabilizing DCLK1.

**Figure 3:**
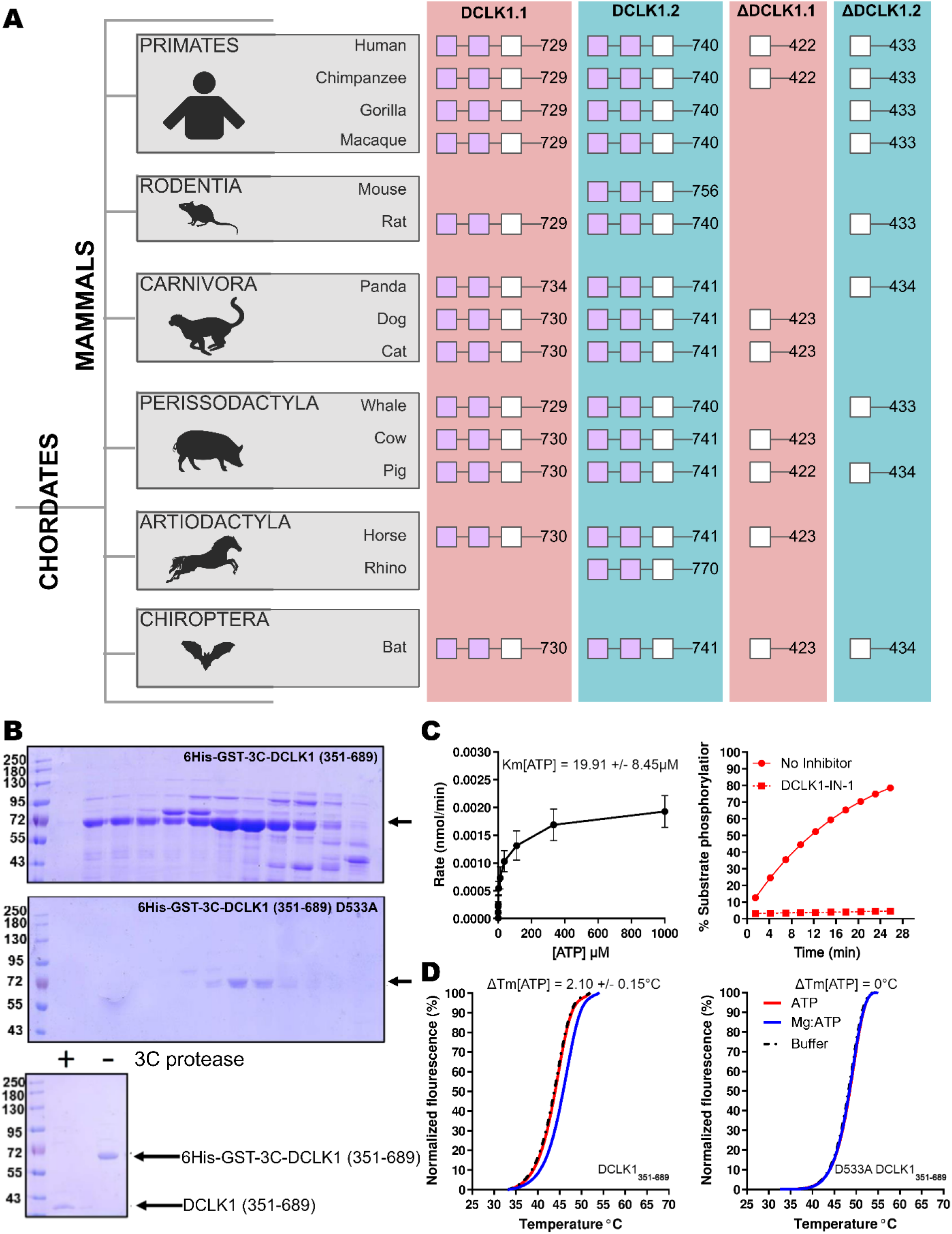
A) Cartoon cladogram of mammalian species showing the domain organization of each DCLK1 isoform from representative annotated sequences from UniProt. UniProt IDs for each sequence are provided in Figure 3**-Source File 1**. **B)** SDS-PAGE of 6His-GST-3C-DCLK1.1 (351-689, Top) or a D533A mutant in which the DFG Asp is mutated to Ala (Middle). Proteins were separated by size exclusion chromatography, and high-purity fractions were pooled. The affinity tag was removed prior to analysis by incubation with 3C protease, leading to a demonstrable shift in mobility (bottom) **C)** Evaluation of catalytic activity towards DCLK1 peptide. DCLK1.1 351-689 possesses a *K*_M[ATP]_ ∼20 μM in vitro (left) and real-time substrate phosphorylation was inhibited by prior incubation with the small molecule DCLK1-IN-1, (right). **D)** Thermal shift assay demonstrating a 2.1°C increase in the stability of DCLK1 351-689 in the presence of Mg:ATP (left), which was absent in the D533A protein (right). Raw data are provided in Figure 3**-Source File 2**.

### Key differences between isoforms in the C-tail of DCLK1 arise from alternative-splicing and different open-reading frames

Higher-order vertebrates have multiple isoforms of DCLK1 and DCLK2, where sequence variations occur in either or both the N and C terminal regions attached to the kinase domain. Human DCLK1, for example, has four unique isoforms. Isoforms 1 and 2 differ in C-terminal tail length due to variations in exon splicing (**Figure 4A**). Further examination of the intron and exon boundaries indicates that human DCLK1.1 contains an additional exon (exon 16) that is not spliced in DCLK1.2. Exon 16 is spliced with exon 17 with a phase 2 intron, which introduces a shift in the reading frame and an earlier translated stop codon (UGA) in exon 17 (**Figure 4B**). In DCLK1.2, exon 15 is spliced with exon 17, with a non-disruptive phase 0 intron, resulting in the full translation of exon 17. These changes introduce multiple indels (insertions and deletions) and result in the insertion of a phosphorylatable threonine (T688) (2) in DCLK1.2 that is absent in the DCLK1.1 variant (**Figure 4B-C**), suggesting a possible exon duplication for adaptive regulation of DCLK1 function by phosphorylation. DCLK1.2 is the best-characterized isoform in terms of structure and function (27), and to compare it with DCLK1.1, we generated a series of C-terminal tail deletion mutants to evaluate how variations in the C-terminal tail contribute to isoform specific DCLK1 functions.

**Figure 4:**
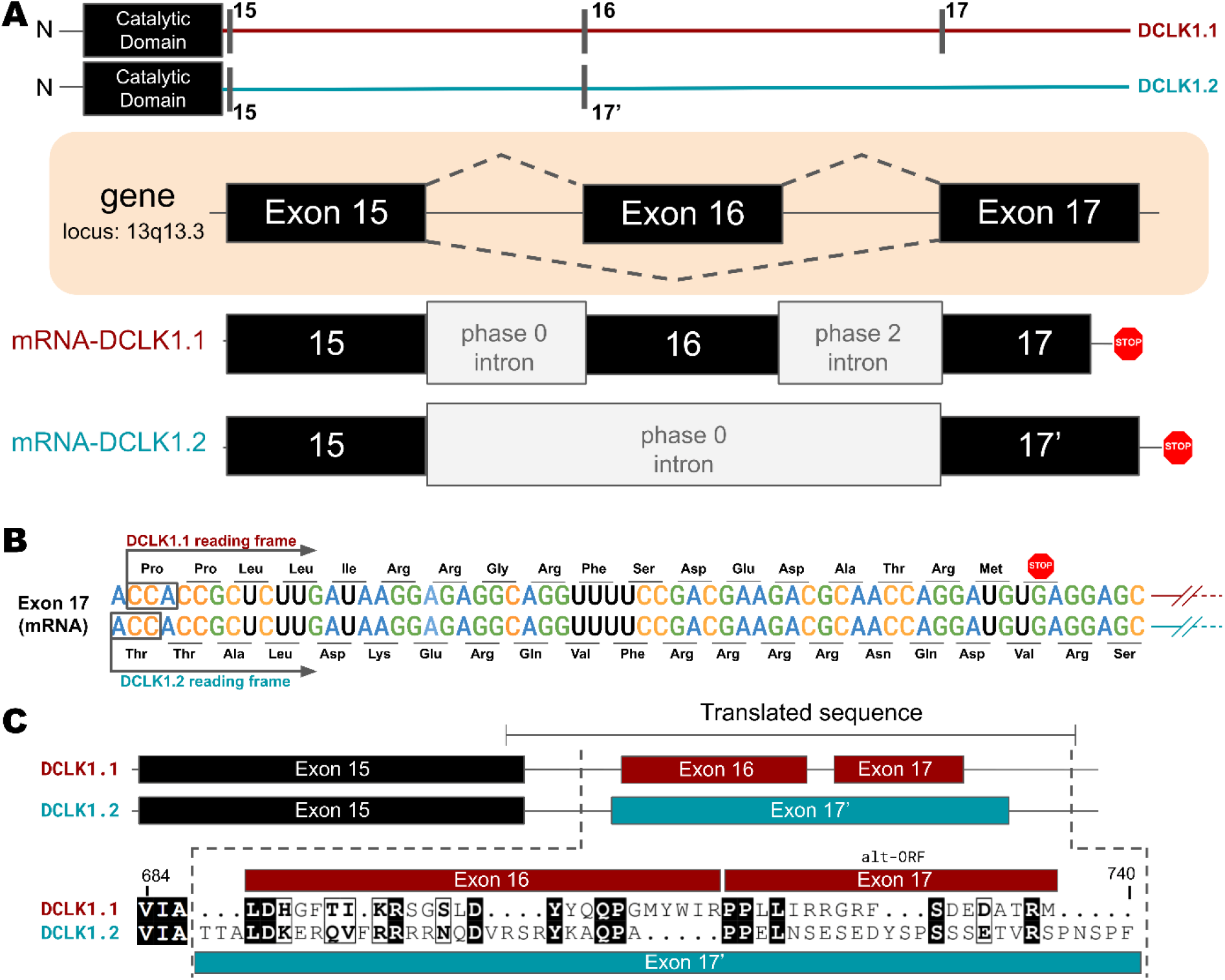
A) Gene and intron-exon organization of DCLK1 human isoforms in the C-terminal tail. The DCLK1 gene is present on locus 13q13.3, and isoforms 1 and 3, contain an additional exon (exon 16), in the C-terminal tail that is absent in DCLK1.2. **B)** A phase 2 intron results in the alternative transcript of exon 17 in isoform 1, translating a different open-reading frame and early stop codon, resulting in the shorter sequence. **C)** Cartoon organization of the C-tail exons (exon 15, 16, and 17) of the DCLK1 isoforms, comparing the translated protein sequence alignment.

### Isoform-specific variations encode changes in molecular dynamics, thermostability and catalytic activity in DCLK1

To study isoform specific differences in the C-tail, we employed experimental techniques to compare protein stability and catalytic activity between purified DCLK1 proteins alongside molecular dynamics simulations for DCLK1.1 and DCLK1.2 with different tail lengths. Isoforms 1.1 and 1.2 share identical sequences across the kinase domain and within the first 38 residues of the C-tail, and we used this information to design a new recombinant protein, termed DCLK1_cat_ (residues 351-686). C-terminal to this totally conserved region, both isoforms possess extended tail segments, which includes the putative inhibitory binding segment (IBS; residues 682-688) and an additional intrinsically-disordered segment (IDS; residues 703-end). To study the role of the C-tail in modulating kinase stability and activity, we purified the DCLK1_cat_, and C-tail containing (long and short) variants of each isoform, each of which lack the N-terminal DCX domains (**Figure 5A**). SDS-PAGE demonstrated that protein preparations were essentially homogenous after affinity and gel filtration chromatography (**Figure 5-figure supplement 1**). The short forms of the recombinant proteins (DCLK1.1_351-703_ and DCLK1.2_351-703_) possess a partially truncated C-tail and were designed to match the amino acid sequence previously used to solve the structure of DCLK1.2 protein (27). Notably, these proteins exclude the IDS. The long forms of the DCLK1 proteins include the full-length C-tail for each isoform (DCLK1.1_351-729_ and DCLK1.2_351-740_) and incorporate IDS domains. We first performed comparative thermal shift analyses to quantify variance in thermal stability between the different purified proteins. When contrasting DCLK1.1_short_ and DCLK1.2_short_ which do not differ in size or tail length but encode unique sets of amino acids in their partially truncated C-tail as a result of alternative splicing (**Figure 4**), we observed that DCLK1.2 was some 14°C more stable than DCLK1.1 (**Figure 5B**). When compared with DCLK1_cat_, both DCLK1.1 short and long exhibited only subtle changes in thermal stability (**Figure 5C & E**), whereas both DCLK1.2 proteins (DCLK1.2_short_ and DCLK1.2_long_) were significantly stabilized (relative ΔTm >16°C, **Figure 5D & E**).

**Figure 5:**
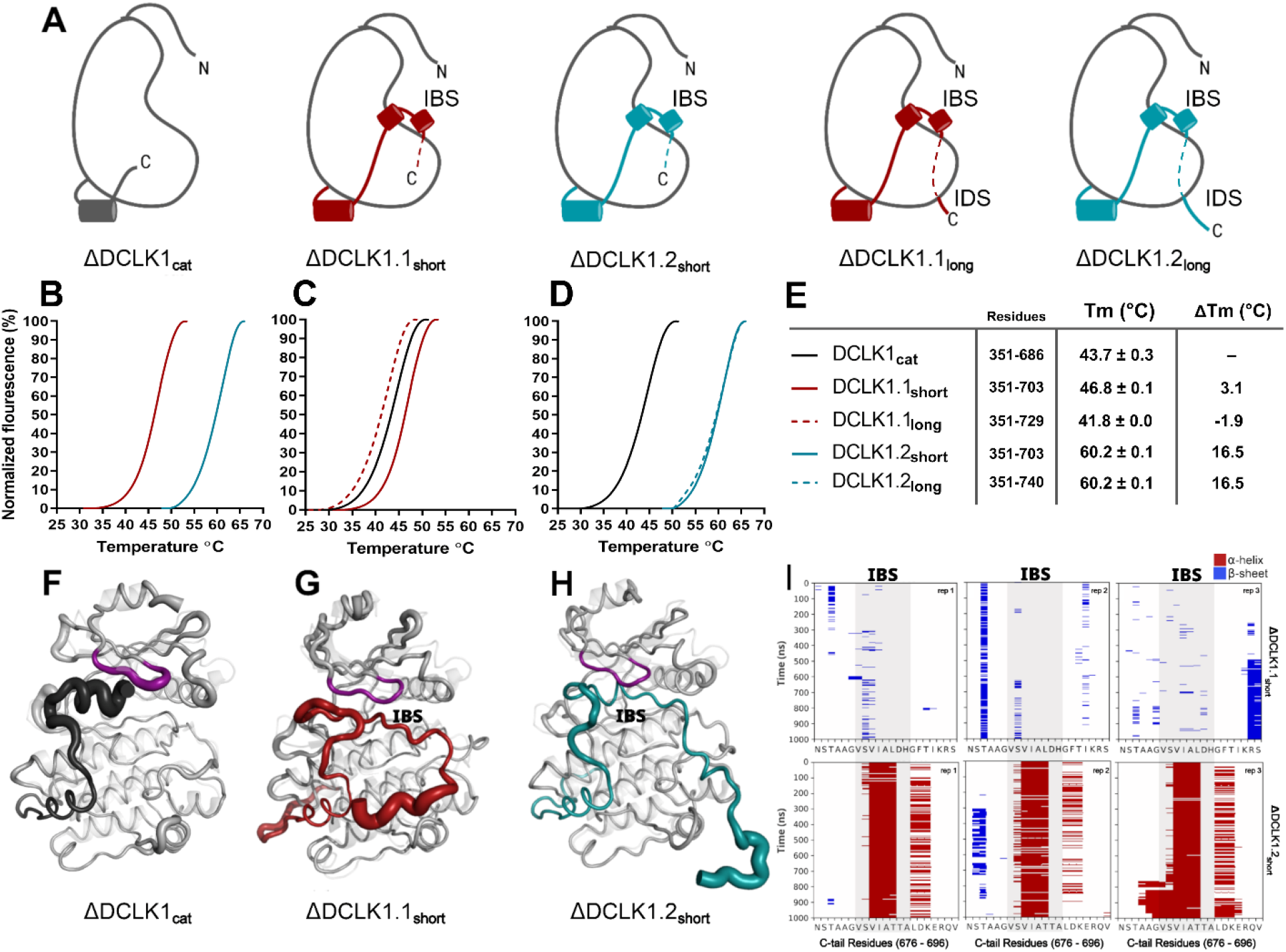
A) Cartoons of DCLK1 construct used in our assays, portraying the locations of the Inhibitory Binding Segment (IBS) and the Intrinsically Disordered Segment (IDS). **B-E)** DSF thermal denaturation profiling of the purified DCLK1 core catalytic domain, or tail-matched DCLK1.1 and DCLK1.2 proteins. Unfolding curves and changes in Tm values (ΔTm) for each protein relative to WT DCLK1_cat_ are indicated. **F-H)** B-factor structural representations of DCLK1_short_ proteins shown in A). The width of the region indicates the extent of flexibility based on averaged RMSF data from three one microsecond MD replicates. **I)** DSSP analysis of three replicates of one microsecond MD simulations showing the residues surrounding the IBS in the C-tail of DCLK1.1_short_ and DCLK1.2_short_. Blue indicates the presence of a Beta-sheet or Beta-bridge secondary structures and red indicates the presence of alpha-helical structures.

We next performed MD simulations to study the dynamics within the distinct DCLK1 C-tails that might explain the observed difference in protein stability. The crystal structure of DCLK1.2 (PDB: 6KYQ) was employed for the DCLK1.2_short_ model and AlphaFold2 was used to model the other proteins (DCLK1_cat_ and DCLK1.1_short_). Comparison of the root mean square fluctuations (RMSF) of the two isoforms in three different replicates of molecular dynamics simulations indicates strikingly different thermal fluctuations in the C-terminal tails and catalytic domains (**Figure 5-source data 1**). In particular, the IBS segment (between 682-688) is stably docked in the ATP binding pocket in DCLK1.2 and an alpha helical conformation is maintained during the microsecond time scale across different replicates (**Figure 5I**, bottom). In contrast, the IBS is more unstable in DCLK1.1, as indicated by high thermal fluctuations and a lack of secondary structure propensity (**Figure 5I**, top). A caveat to bear in mind is that DCLK1.1 is an AlphaFold2 model, which will also account for increased RMSF. Analysis of sequence variations and structural interactions provides additional insights into the differential dynamics of the two isoforms. The helical conformation of the IDS in DCK1.2 is maintained during the simulation due, in part, to a capping interaction with Thr 687, which is absent in DCLK1.1 due to the alternative splicing event detailed above. Likewise, another key residue in DCLK1.2, Lys 692, anchors the tail to the catalytic domain through directional salt bridges with the conserved aspartates (Asp 511 and Asp 533) in the HRD and DFG motifs (**Figure 5, figure-supplement 2A**). These interactions are not observed in DCLK1.1 simulations because Lys 692 is substituted to a histidine (His 689), which is unable to form a corresponding interaction with the catalytic domain (**Figure 5, figure-supplement 2B**). We also evaluated the effects of T688A (non-phosphorylated) or T688E (phosphomimetic) mutations through DCLK1 MD simulations and found that the two mutations slightly destabilize the tail relative to WT. Three replicates of the two mutants show increased RMSF of the tail region relative to WT DCLK1.1 (**Figure 5, figure-supplement 3**). Either mutation was not sufficiently destabilizing on its own to unlatch the C-tail, and we hypothesize that other residues in addition to T688 are also likely to be important for contributing to conformational regulation of the kinase domain by the C-terminal tail. The variable docking of the C-tail within the kinase domains of the two DCLK1 isoforms, and the extent to which this contributes to more transient or stable autoinhibited states are explored in more detail in the next section.

### Residues contributing to the co-evolution and unique tethering of the C-terminal tail to the DCLK catalytic domain

To identify specific residues that contribute to the unique modes of DCLK regulation by the C-terminal tail, we performed statistical analysis of the evolutionary constraints acting on DCLK and related CAMK family sequences. We aligned the catalytic domain of DCLK and related CAMK sequences from diverse organisms and employed the Bayesian Partitioning with Pattern Selection (BPPS) method (30) to identify residues that most distinguish DCLK sequences (foreground alignment in **Figure 6B**) from CAMK sequences (background alignment). Beyond the catalytic domain, DCLKs share sequence and structural similarities in the first helix of the tail (αR1 in CAMK1) (31,32), with other CAMKs but share no detectable sequence similarity beyond this helical segment. DCLKs also share a CAMK-specific insert segment located between F and G helices in the catalytic domain, although the nature of residues conserved within the insert is unique to individual CAMK families (**Figure 6-figure supplement 1**). BPPS analysis revealed DCLK-specific constraints in different regions of the kinase domain, most notably, the ATP binding G-loop, N terminus of the C-helix, the activation loop, and C-terminus of the F-helix (**Figure 6-figure supplement 2**).

**Figure 6:**
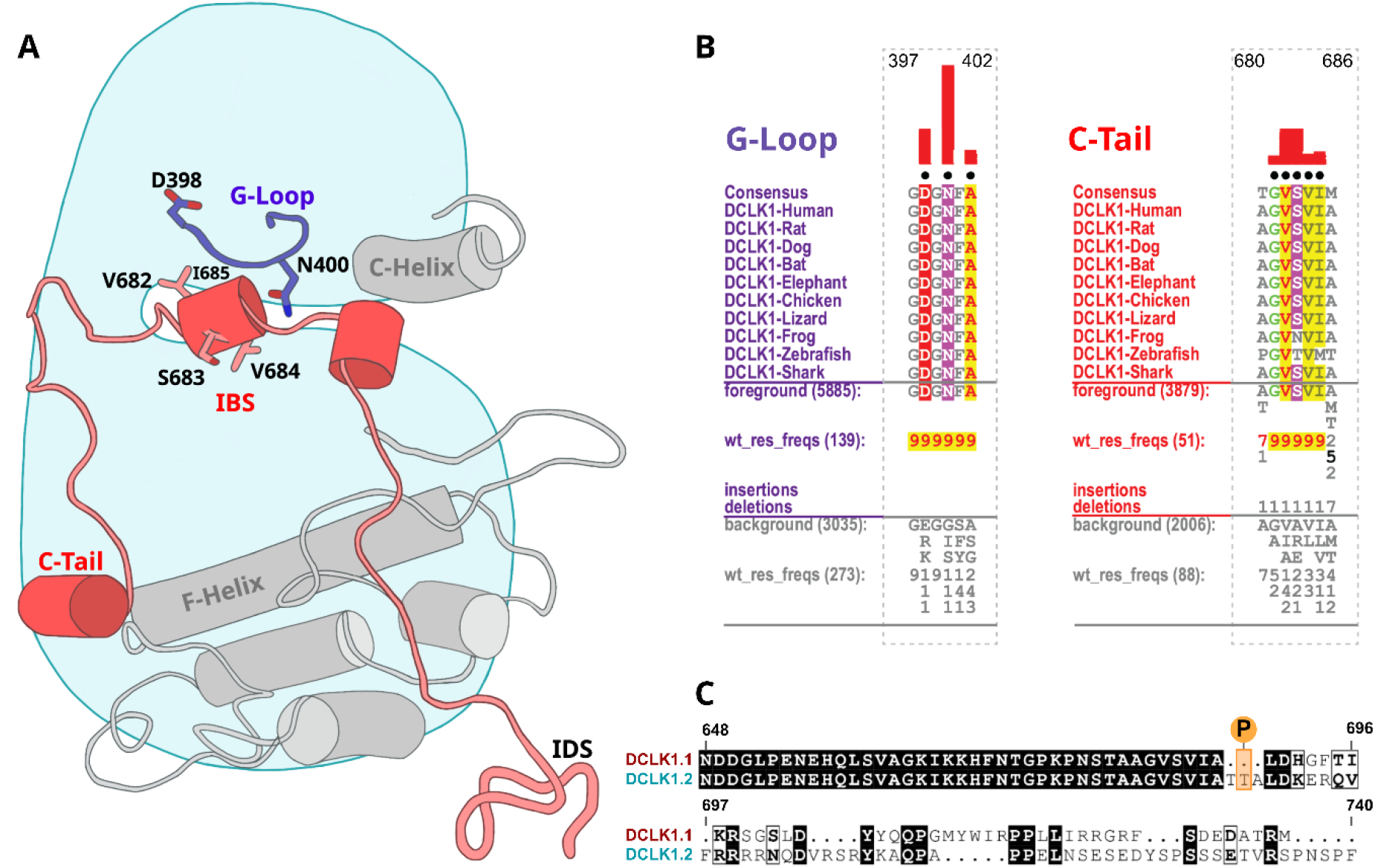
Identification of DCLK specific constraints. A) Cartoon of DCLK1.2 and the intrinsically disordered segment (IDS) with evolutionary constraints mapped to the kinase domain and C-tail. **B)** Sequence constraints that distinguish DCLK1/2/3 sequences from closely related CAMK sequences are shown in a contrast hierarchical alignment (CHA). The CHA shows DCLK1/2/3 sequences from diverse organisms as the display alignment. The foreground consists of DCLK sequences while the background alignment contains related CAMK sequences. The foreground and background alignments are shown as residue frequencies below the display alignment in integer tenths (1–9). The histogram (red) indicates the extent to which distinguishing residues in the foreground diverge from the corresponding position in the background alignment. Black dots indicate the alignment positions used by the BPPS (Neuwald, 2014) procedure when classifying DCLK sequences from related CAMK sequences. Alignment number is based on the human DCLK1.2 sequence (UniProt ID: O15075-2). **C)** Sequence alignment of human DCLK1 isoforms.

Some of the most significant DCLK specific residue constraints map to the ATP binding G-loop (GDGNFA motif) (Figure 6A-B). In particular, Asp 398, Asn 400 and Ala 402 are unique to DCLKs as the corresponding residues are strikingly different in other CAMKs. Asp 398 is typically a charged residue (K/R) in other CAMKs while Asn 400 and Ala 402 are typically hydrophobic and polar residues, respectively (see residue frequencies in background alignment; **Figure 6B**). Notably, both Asn 400 and Asp 398 make direct interactions with residues in the C-tail either in the crystal structure or molecular dynamics simulations (see below). Likewise, DCLK conserved residues in the C-helix and activation loop tether the C-terminal tail to functional regions of the kinase core, suggesting co-option of the DCLK catalytic domain to uniquely interact with the flanking cis regulatory tail.

### An autoinhibitory ATP-mimic completes the C-spine and mimics the gamma phosphate of ATP

The most stable segment of the C-tail based on the B-factor and RMSF fluctuations in MD simulations is a unique region (682-688 in DCLK1.2) that docks into the ATP binding pocket through both hydrophobic and hydrogen-bonding interactions. Remarkably, this peptide segment mimics the physiological ATP ligand, and stabilizes the catalytic domain through ‘completion’ of the hydrophobic catalytic spine (**Figure 7A**, (33)). The residues that mimic adenosine and complete the C-spine of DCLK1 are Val 682, Val 684, and Ile 685 (PDB: 6KYQ), which are part of the α-helix that docks to the ATP-binding pocket (**Figure 7B**). Interestingly, based on our BPPS analyses, these C-tail residues are uniquely vertebrate DCLK1-specific pattern constraints. At the tail end of this α-helix are two Thr residues, Thr 687 and Thr 688. As previously noted, these Thr residues mark the beginning of exon 17, and are one of the key variations between human DCLK1 isoforms, found only in DCLK1.2 variants.

**Figure 7:**
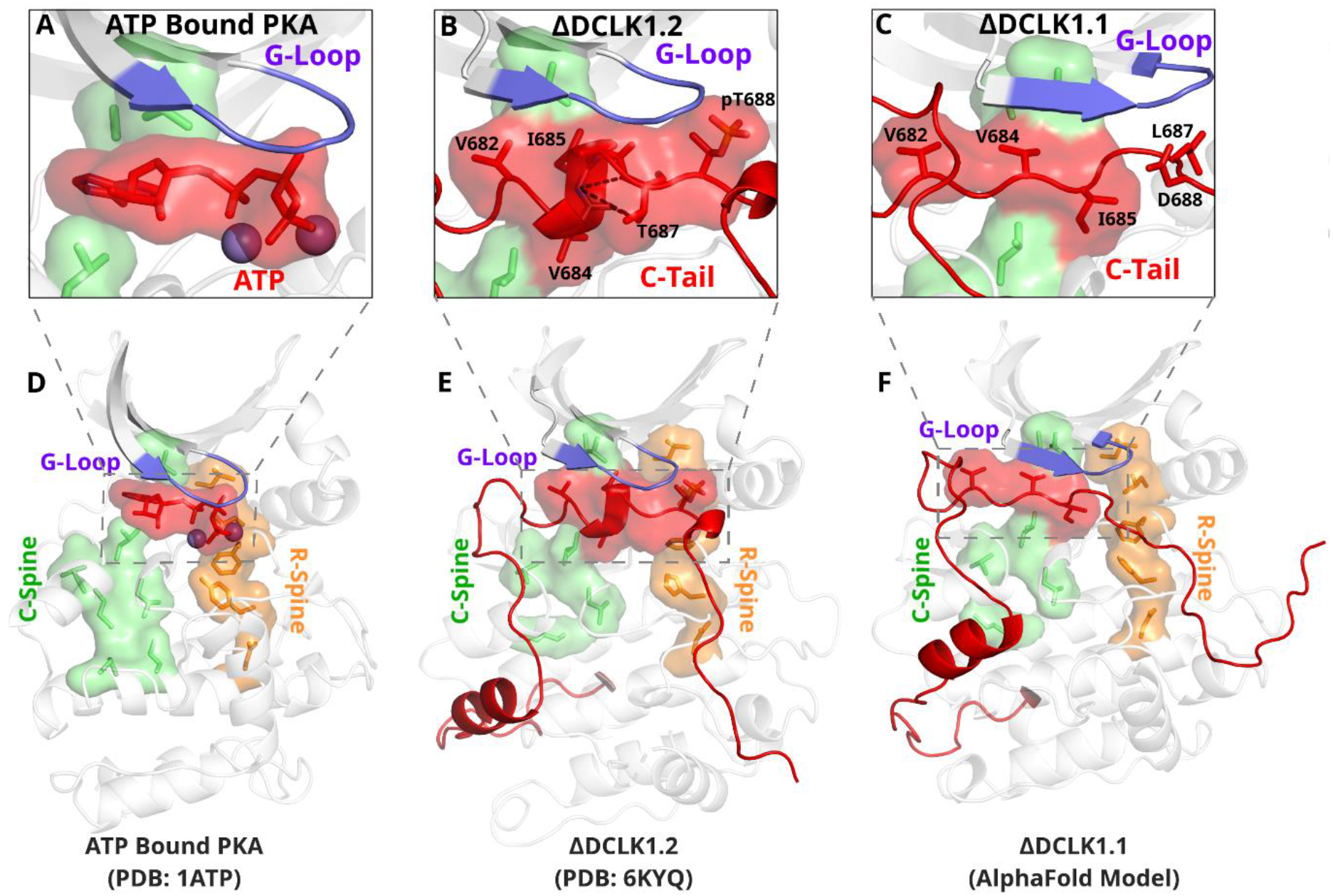
The DCLK1 C-tail ‘completes’ the regulatory C-spine (green). A) PKA crystal structure (PDB: 1ATP) with bound ATP in red and Mg^2+^ in purple. The C-spine is completed by the adenine ring of ATP. The gamma phosphate of ATP hydrogen bonds with the second glycine of the G-loop. **B)** DCLK1.2 crystal structure (PDB: 6KYQ) showing how the C-tail (red) docks underneath the pocket and mimics the ATP structure. The C-spine is completed by V682 and V684 in the C-tail and helical segments defined using DSSP are shown. T687 is also depicted making multiple hydrogen bonds with the backbone of V684 and I685 (dashed lines). **C)** DCLK1.1 AlphaFold2 model showing an unstructured loop in the C-tail docking into the ATP binding pocket, where V684 and I685 are predicted to complete the C-spine. The average per-residue confidence of the C-tail is 49%. **D-F)** Zoomed out versions of A-C, demonstrating how the DCLK1 C-tail docks into the ATP binding cleft, akin to ATP in PKA.

Structural analysis and MD simulations reveal that Thr 687 in DCLK1.2 caps the stable α-helix that extends the C-spine (**Figure 7B, Figure 7-figure supplement 1B**). In comparison, the same region in DCLK1.1, which lacks Thr 687, is predicted to be unstructured. Upon phosphorylation, Thr 688 in DCLK1.2 can mimic the gamma phosphate of ATP by maintaining a stable hydrogen bonding distance with the backbone of the second glycine of the G-loop (G399) (**Figure 7B, Figure 7-figure supplement 1C**). We additionally observe that the sidechain of Asn 400, a DCLK-specific G-loop constraint, further stabilizes the phosphate group in pThr 688 through hydrogen bonding. As previously described, Thr 688 is unique to DCLK1.2. The lack of this functional site in DCK1.1 is correlated with increased RMSF and instability of the ATP-mimic segment in isoform 1 MDs (**Figure 5C-D, Figure 7-figure supplement 1B**). Comparatively, MD analysis of DCLK1.2 and a phosphothreonine-containing DCLK1.2 demonstrates reduced C-tail fluctuations, suggesting the potential regulatory involvement of Thr 688 phosphorylation for further modulation of the autoinhibited conformation (**Figure 7-figure supplement 1C**), consistent with previous findings (2).

### Mutational analysis support isoform-specific allosteric control of catalytic activity by the C-terminal tail

To evaluate how sequence differences between DCLK1.1 and DCLK1.2 affected both thermal stability and catalytic potential, we generated targeted mutations at contact residues within the Gly-rich loop and C-tail of DCK1.1 and DCLK1.2 (at the indicted residues depicted in **Figure 8A**) which we predicted would disrupt or destabilize C-tail docking within the domain. All proteins were purified to near homogeneity by IMAC and size exclusion chromatography (**Figure 5-figure supplement 1**), and the thermal stability of a panel of DCLK1.2 mutant and WT proteins were compared side-by-side with the DCLK1_cat_ (**Figure 8B**). The ΔTm values obtained (**Figure 8C**) demonstrate that mutation of Asp 398 or Asn 400 in the Gly-rich loop are by themselves insufficient to destabilize DCLK1.2. In marked contrast, dual mutation of the hydrophobic pair of Val 682 and Val 684 residues to Thr, or mutation of the acidic tail residue Asp 691, resulted in a pronounced reduction in DCLK1.2 thermal stability. Moreover, the recorded Tm values for these latter two mutations quite closely resembled the Tm of DCLK1_cat_ (which lacks the C-tail entirely), which is consistent with the uncoupling of the C-tail and a commensurate decrease in thermal stability associated with loss of this interaction. We next determined the catalytic activity of our recombinant DCLK1.1 and DCLK1.2 proteins side-by-side (**Figure 8D, Figure 8-figure supplement 1**). Although partially diminished in relation to DCLK1_cat_, both DCLK1.1_short_ (351–703) and DCLK1.1_long_ (351–729) variants possess robust catalytic activity. This suggested ineffective ATP-competitive auto-inhibition mediated by the C-tail segment of DCLK1.1 and is consistent with their closely matched Tm values to DCLK1_cat_ (**Figure 5C**). Interestingly, both C-tail containing variants of DCLK1.1 (and particularly DCLK1.1^351-729^) exhibited lower affinity for ATP (inferred from *K*_M[ATP]_ for peptide phosphorylation), which is consistent with partial-occlusion of the ATP binding pocket (**Figure 8-figure supplement 1**). In marked contrast, the detectable kinase activity for short (351–703) or long (351–740) DCLK1.2 proteins was significantly blunted compared to DCLK1_cat_, exhibiting just ∼5% of the activity of the catalytic domain alone, and consistently, the calculated *K*_M[ATP]_ was ∼ 4 fold higher compared to the catalytic domain lacking the C-tail. We also utilized autophosphorylation as a proxy for overall kinase activity. Quantitative tandem mass spectrometry (MS/MS) analysis of site-specific autophosphorylation within DCLK1.1_short_ and DCLK1.2_short_ demonstrate a marked reduction in the site-specific abundance of phosphate in DCLK1.2 when compared to DCLK1.1 at two separate sites that could be directly and accurately quantified by MS (S438 and S660, DCLK1.1 relative abundance set to 1, **Figure 8-figure supplement 2**). LC-MS/MS also indicated that several autophosphorylation sites identified in isoform 1 were absent in DCLK1.2 (Ser 683 and Thr 692, the latter of which is an amino acid that is unique to the C-tail of DCLK1.1, **Figure 8-figure supplement 2**). Interestingly, amino acid substitutions in the G loop or the C-tail of DCLK1.2 designed to subvert C-tail and ATP site interactions also had major effects on DCLK1.2 phosphorylation and catalytic activity. DCLK1.2 D398A was activated some 5-fold when compared to the WT form, whereas DCLK1.2 N400A was almost as active as the DCLK1_cat_. Consistently, DCLK1.2 V682T/V684T and D691A proteins were also much more active than the WT form of DCLK1.2. Kinetic analysis also confirmed higher *Vmax* (but broadly similar *K*_M[ATP]_) values for DCLK1.2 D398A and V682T/V684T relative to the WT protein (**Figure 8-figure supplement 1**). Moreover, comprehensive LC-MS/MS phosphosite mapping revealed a marked increase in the total number of phosphorylated amino acids in all of the mutant DCLK1.2 proteins, consistent with the enhanced catalytic activity of these proteins when compared to WT DCLK1.2 (**Figure 8-figure supplement 2**). Together, these observations confirm that targeted mutations are sufficient to relieve ATP-competitive C-tail autoinhibition by physical tail uncoupling; this model is also strongly supported by marked changes in the biophysical stability of the mutant proteins, particularly for V682/684T and D691A (**Figure 8B**).

**Figure 8:**
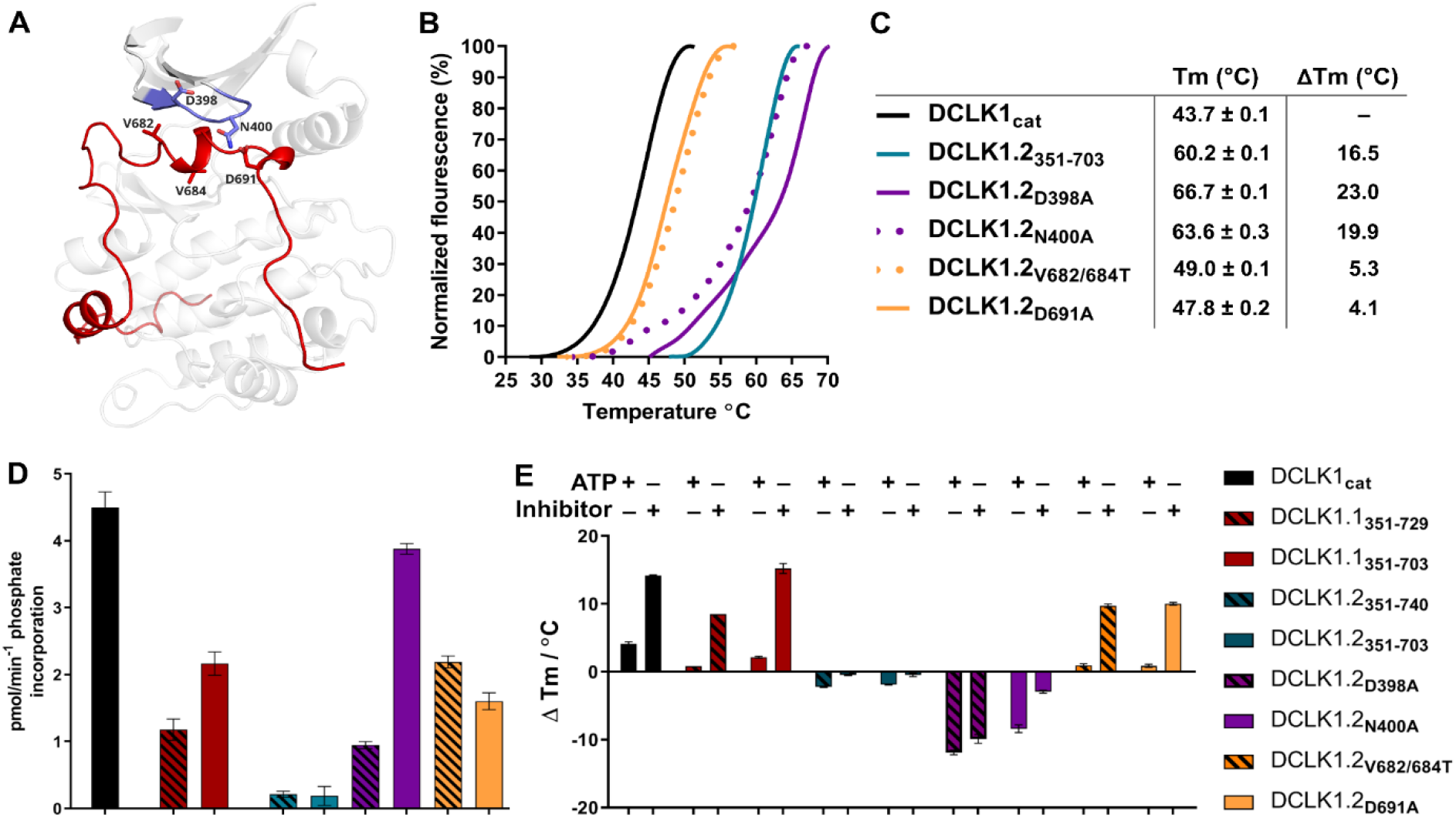
A) Structural depiction of DCLK1.2 (PDB: 6KYQ) showing the location of modified DCLK1 amino acids on the G-loop (purple) or C-tail (red). **B-C)** Differential Scanning Fluorimetry assays depicting thermal denaturation profiles of each protein along with the calculated Tm value. **D)** Kinase assays. DCLK1-dependent phosphate incorporation (pmol/min_-1_) into the DCLK1 peptide substrate was calculated for DCLK1_cat_, long and short DCLK1.1 and the indicated DCLK1.2 variants. **E)** Thermal stability analysis in the presence of ATP or DCLK1-IN-1 for DCLK1 proteins. For DCLK1.2, all proteins were generated in the DCLK1.2 short background.

To expand on this finding, we investigated the interaction of Mg:ATP or DCLK1-IN-1 to our panel of DCLK1.1 and 1.2 proteins (**Figure 8E**), using changes in thermal stability as a reporter of ligand binding. DCLK1_cat_ (351–686), DCLK1.1_short_ (351–703) and DCLK1.1_long_ (351–729) proteins all behaved similarly in the presence of either Mg:ATP or DCLK1-IN-1, inducing marked stabilization. In contrast, DCLK1.2_short_ (351–703) or DCLK1.2_long_ (351–740) proteins registered negligible thermal shifts in the presence of the same concentration of either ligand, which is in-line with the C-tail tightly occupying the ATP-binding site and obstructing their binding. Remarkably, D398A and N400A DCLK1.2, whose high basal Tm values (compared to DCLK1_cat_) are consistent with stabilization by docking of the C-tail within the kinase domain, were markedly destabilized in the presence of either ligand. This suggests that incorporation of these G-loop mutations in isolation is insufficient to dislodge the C-tail, but rather that the stability of the interaction is compromised to the extent that either ATP or DCLK-IN-1 can competitively dislodge the bound tail from the ATP active site, resulting in a net destabilization caused by lack of tail engagement. This observation is corroborated by the results of our kinase assays, where both D398A and N400A mutants were more active than WT DCLK1.2, confirming appropriate ATP binding (a pre-requisite for catalysis). Finally, DCLK1.2 V682T/V684T and D691A, which exhibit lower basal Tm values than the WT protein (indicating a loss of tail interaction), were both stabilized in a ligand-dependent manner to a similar degree to that observed for DCLK1_cat_ and DCLK1.1 proteins (**Figure 8E**). Collectively, these observations clearly demonstrate that the C-tail section of DCLK1.2 can both stabilize the canonical DCLK kinase domain and inhibit kinase activity (by impeding the binding and structural coordination of Mg:ATP) much more effectively than that of DCLK1.1. Our targeted mutational analysis of key contact residues in DCLK1.2 also clearly shows that this is a consequence of specific amino acid interactions that are absent in DCLK1.1 due to alternative-splicing and subsequent sequence variation.

### Classification of DCLK regulatory segments

We synthesized our experimental findings by classifying the DCLK C-tail into six functional segments, based on interactions with different regions of the catalytic domain and their conservation between the C-tail splice variants in our analysis (**Figure 9**): First is an ATP-mimetic peptide segment (residues 682-688 in DCLK1.2) that readily mimics physiological ATP binding by completing the C-spine in the nucleotide-binding site. The inhibitory peptide also contains a phosphorylatable Thr residue, which sits adjacent to the highly characteristic Gly-rich loop (GDGNFA, residues 396-402). Second, we define a pseudosubstrate mimic (PSM, residues 692-701), which interacts with the acidic HRD and DFG Asp side-chains and docks in the substrate pocket occluding substrate access. Third, at the C-terminus of the tail, lies an intrinsically disordered segment (IDS, residues 702-749, **Figure 9-figure supplement 1**), which packs dynamically against DCLK conserved residues in the kinase activation loop. Fourth, at the beginning of the C-tail lies a CAMK-tether (residues 654-664), a set of residues that pack against a CAMK-specific insert in the C-lobe. In many CAMK crystal structures, this insert makes multiple contacts with the F-helix and C-tail (**Figure 6-figure supplement 1**). Fifth, this is followed by a highly dynamic pseudosubstrate region (residues 672-678) that occludes the substrate pocket and will thus interfere with substrate phosphorylation. Sixth, a transient beta-strand is formed in DCLK1.2 through amino acid specific sequences that help modulate and potentially strengthen binding of the C-tail in this isoform (**Figure 7-figure supplement 1A-B**). Collectively, these segments and their associated interaction sites demonstrate that co-evolution of the unique C-tail with the catalytic domain is the central hallmark of DCLK functional divergence, and that changes in these segments possess the ability to ‘supercharge’ catalytic output of the kinase. In particular, the variable C-terminal segments of the tail might contribute to isoform-specific functional specialization. The combinatorial diversity of events that modulate C-tail function may allow DCLKs to nimbly coordinate various tasks including ATP-binding, substrate-based phosphorylation, regulation of DCX domain phosphorylation and structural disposition, kinase autoinhibition and allosteric regulation. Isoform-specific variability provides additional nuance to regulatory and catalytic signaling events and may even contribute to differences in cellular localization (e.g. cytoplasm or nucleoplasm) and tissue-specific activity, enabling contextual DCLK regulation through these modular sequence segments.

**Figure 9:**
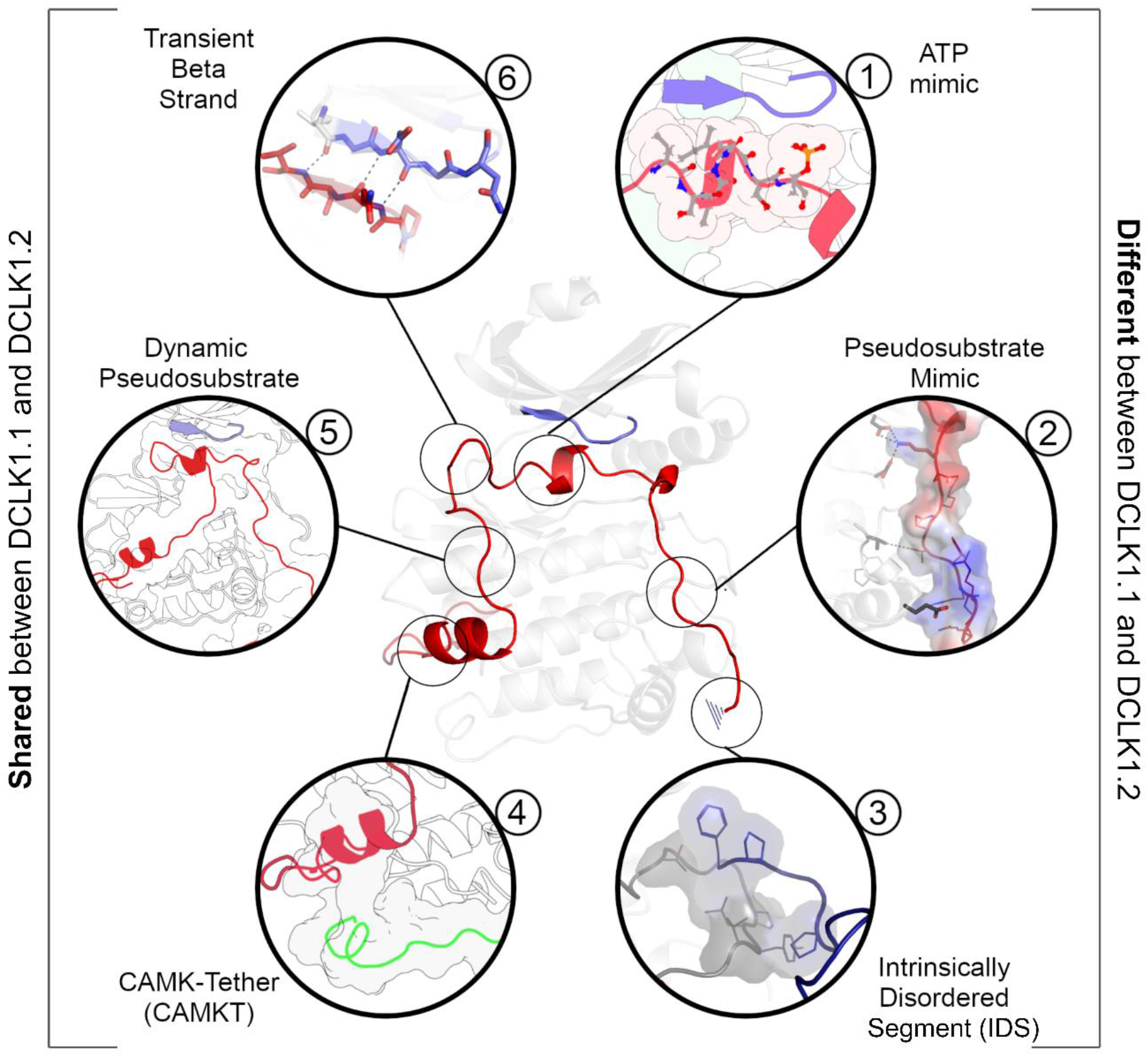
A DCLK1 C-tail can act as a multi-functional Swiss Army Knife, using six distinct segments for a variety of regulatory functions including mimicking ATP binding/association, stabilizing the G-loop, occluding the substrate binding pocket, and packing against the kinase activation loop.

## Discussion

The kinase domain is a conserved switch for phosphorylation-based catalytic regulation. Yet the complexity of cell signaling pathways demands other nuanced forms of regulation beyond binary “on” or “off” switch-based mechanisms. For many Ser/Thr kinases, including AGC and CAMK families, these distinct regulatory functions come from segments which flank the kinase domain, N– and C-terminal regions, which serve to modulate activity through allosteric activation, inhibition, or rheostatic behaviors that change based on environmental conditions (4). In this study, we expand on our knowledge of allosteric diversity in the human kinome by revealing how alternative splicing of the DCLK1 C-tail contributes to isoform-specific behaviors, coupling regulation of catalytic output, phosphorylation, protein dynamics and stability, substrate binding, and protein-protein interactions. Our “Swiss Army Knife” model for DCLK1 expands our view of allosteric regulation as not just a dynamic process facilitated by proteins, but one where adaptive genetic mechanisms, like differential splicing, dexterously tune isoform-specific functions for specific cellular signaling roles; in the case of DCLK1.1, this allows ‘supercharging’ of catalysis between splice variants due to key amino acid differences in the C-tail that are lacking in the DCLK1.2 isoform.

Multiple members of the human kinome have independently evolved C-tail regions that dock to the N or C-lobe of the kinase domain *in cis*. In the case of the AGC kinases, the C-terminal tail is a very well-studied *in-cis* modulatory element that serves to explain a variety of regulatory properties in this kinase sub-family (14,34–36). Classical deletion studies with members of the CAMK family, have also revealed a cis-acting inhibitory element lying C-terminal to the catalytic domain of both CAMK1 (37) and CAMK2 (38). More recent examples of C-tail functional diversity in CAMK family members are presented by the human pseudokinases TRIB1 and TRIB2, which employ C-tail sequences to either latch (and structurally restrict) the atypical kinase domain or to bind competitively to the Ubiquitin E3 ligase COP1 (39,40). Functional disengagement of the TRIB1 or TRIB2 tail through deletion, mutagenesis or small molecule binding has marked effects on pseudokinase conformation, intrinsic stability and cellular transformation (41–44).

### A novel pseudosubstrate region encoded by DCLK1

In addition to the marked differences between DCLK1 splice variants relevant to nucleotide binding, small molecule interactions and catalysis, our work also reveals two unique pseudosubstrate segments present before and after the IBS. Before the IBS segment, we observe the formation of an anti-parallel transient beta sheet with the beta1 strand in the catalytic domain (**Figure 5I, Figure 7-figure supplement 1A-B**). During the formation of this transient structure, the C-tail dynamically occludes part of the substrate binding pocket. A beta-sheet is observed in all three MD replicates of DCLK1.1, but only in a single replicate of DCLK1.2 dynamic analysis. At the other end of the IBS is another pseudosubstrate segment whose structure and dynamics change as a result of alternative exon splicing. In DCLK1.2, the pseudosubstrate segment is stable, with an average RMSF of 1.8 Angstroms, facilitated by key interactions from Lys 692, which coordinates acidic residues in the HRD and DFG motifs (**Figure 5-figure supplement 2A**). In DCLK1.1, Lys 692 is replaced by a His, which weakly coordinates with the HRD and DFG motifs, resulting increased dynamics of the segment and an RMSF of 3.1Å (**Figure 5, Figure 5-figure supplement 2B**). Together, residue variation between isoforms contribute to differences in stability and alteration of dynamics of the tail. HPCAL1 was recently proposed as a possible substrate that activates DCLK1 in a calcium-dependent manner, but it is unclear how it may bind DCLK1 (26). Because only exon 15 of the C-tail is conserved between the isoforms, it is possible the location of binding occurs in this dynamic pseudosubstrate segment prior to the IBS, where increased flexibility and occlusion of the substrate pocket is reflective of the absence of HPCAL1, or a similar calmodulin-like substrate.

### Discovery of the DCLK1 ATP-Mimic region; a splice-variant specific regulatory module

Our structural analysis of DCLK1 reveals a remarkable structural mimic of ATP located in the C-tail, which differs markedly between DCLK1.1 and DLCK1.2 splice variants. We note for the first time that a set of three residues in the ATP-mimic, Val 682, Val 684, and Ile 685, are conserved across all isoforms of DCLK1 and DCLK2 (Figure 1C) and serve to extend the kinase C-spine. Mutation of these residues in DCLK1.2 uncouples tail binding and activates the kinase. Proximal to these residues are two Thr residues (Thr 687 and Thr 688), which are present in DCLK1.2, but absent in DCLK1.1. Based on published phosphoproteomics data, both Thr residues can be phosphorylated (2) and are thus likely to be regulatory in DCLK1.2. Although we could not detect phosphorylation of either of these predicted regulatory sites in the WT form of DCLK1.2, we consistently observed pThr 688 in activated mutant DCLK1.2 variants (**Figure 8-figure supplement 2**). By analyzing DCLK1 dynamics using MD simulations, we observed multiple key interactions between the G-loop and the C-tail in DCLK1, such as dipole interactions with the second glycine in the G-loop by the phosphothreonine. In addition, Thr 687 contributes to increased stabilization of DCLK1.2 tail by forming a C-cap with the helical ATP-mimic segment. We aligned the intensively-studied protein kinase (PKA, PDB: 1ATP), a DCLK1.2 structure (PDB: 6KYQ), and frames of our MD trajectory, which demonstrate remarkable overlap of the ATP gamma phosphate and C-tail phosphothreonine, which seemingly acts as a mimic for the ATP co-factor. As phosphorylation is reported to lead to DCLK1 inhibition, this suggests a complex mechanism of regulation, in which the DCLK-specific constraints in the G-loop, the intrinsic flexibility of the C-tail, and threonine phosphorylation, by cis or trans-mediated modification, systematically prevent hyperphosphorylation of the doublecortin domains and cellular effects. Somewhat paradoxically, we could only identify phosphorylated Thr 688 in activated DCLK1.2 mutants, but not in the autoinhibited (WT) versioin. form. This suggests that the selected mutations exhibit a regulatory hierarchal dominance over inhibitory Thr 688 phosphorylation and are sufficient to liberate DCLK1.2 from its auto-inhibited, C-tail bound state. This also implies that phosphorylation of Thr 688 may only be minimally-required for autoinhibition, especially given its association with the hyperactive variants obtained by mutagenesis (**Figure 8-figure supplement 1-2)**.

Finally, we have evaluated the terminal residues in the DCLK1 C-tail, which are predicted to be intrinsically disordered. Side-by-side analysis of DCLK1.1 and DCLK1.2 in which this region is added from a common core terminating at residue 703, shows that increasing the length of the tail in both DCLK1.1 and DCLK1.2 has little additional effect on the inhibitory or stabilizing effects driven by the highly ordered tail regions that precede it. Kinase domains are regulated by IDRs in a multitude of ways, but the CAMK family is specifically enriched for adapted C-terminal extensions that, as we show here, can block the ATP and substrate binding, and enzymatically inactivate the kinase domain by occlusion of the activation loop through a flexible helical IDS on their C-tail (4). We note that the DCLK1.2 kinase domain crystallizes in an ‘active’ closed conformation, despite binding of the C-tail in an autoinhibitory manner (27). Repeated packing motions of the IDS against the activation loop in all replicates of DCLK1.1 MD simulations, suggest that tail may occlude the activation loop, similar to other CAMKs, possibly pointing to a mode of *cis* autoregulation. Conversely, AlphaFold2 predicts the placement of the DCX domains as adjacent to the IDS in both DCLK1.1 and DCLK1.2 (Figure S1). It is possible, like other CAMKs, the IDS facilitates protein binding, whether to the DCX domains, calcium-modulated proteins, or other kinases.

### Evolutionary divergence and functional specialization of DCLKs

For the DCLK family as a whole, we discovered phylogenetic divergence between DCLK1 and 2 as a relatively recent event, (Figure 1A) in which metazoan DCLK3 is the more ancestral DCLK gene from which DCLK1 and 2 emerged after duplication. Because of shared evolutionary constraints and the recent divergence between DCLK1 and 2, we surmise the functional specialization of the DCLK1 tail is shared between these paralogs. Moreover, we quantify key differences between human DCLK1.1 and 1.2 activity that are impacted by amino acid changes that contribute to the function of the C-tail. The differences between DCLK1 isoforms 1 and 2 are generated by variations in exon splicing, which change both the C-tail protein sequence, and introduce or exclude potential phosphorylation sites. Expression of the highly autoinhibitable DCLK1.2 isoform is believed to be predominant in the brain during embryogenesis, although DCLK1.1 is also thought to be present in the adult brain (45). It is therefore possible that an altered ratio in DCLK1.1/1.2 expression, accompanied by changes in the requirement for DCLK1 auto-regulation, are relevant for early neurogenesis. The overall sequence similarity at the protein level, despite the loss of a pair of putative phosphorylation sites, suggests a possible exon duplication in the C-tail, whereby polymorphisms have allowed for adaptive regulation during development and proliferation. Indeed, we also speculate that the induced expression of DCLK1.2 splice variants in multiple cancer subtypes (46) is likely to be indicative of a survival and drug-resistance role that could be targetable with new types of small molecule. Although nanomolar DCLK1 ATP-site inhibitors such as DCLK1-IN-1 have been developed that can bind tightly to the DCLK1 ATP site (**Figure 3** and **Figure 3-figure supplement 1**), the ‘problematic’ existence of human DCLK1.1 and DCLK1.2 splice variants with distinct auto-inhibitory properties may present a challenge to compound engagement in the cell, where relief of auto-inhibition through C-tail undocking in DCLK1.2 is likely to require a high concentration of compound in order to compete and disengage interactions at the ATP site. Indeed, although potent chemical DCLK1 inhibitors such as DCLK1-IN-1 are known to influence DCLK1 autophosphorylation and cell motility, they have relatively modest effects in cells in terms of cytotoxicity (29,47). Therefore, we propose that the dual inhibitory effects of the C-tail and the transmission of this information to adjacent DCX domains, which control adaptive cellular phenotypes such as EMT in cancer cells (48), may make allosteric classes of DCLK1 inhibitor a preferred therapeutic option, especially if they can be tailored specifically towards DCLK1.1 or DCLK1.2, whose autoregulation is different in terms of the varied molecular details we have uncovered here.

## Materials and Methods

### Ortholog Identification

To identify orthologs, we used the software KinOrtho (49) to query one-to-one orthologous relationships for DCLK1/2/3 across the proteome. After collation of the various orthologs, we parsed the sequence data for taxonomic information and classified each sequence by family. We further separated human DCLK1 into each unique isoform and aligned them.

### Phylogenetic Analysis

We identified diverse DCLK orthologs from the UniProt database (50) using an profile-based approach (51). From this dataset, we manually curated a taxonomically diverse set of DCLK orthologs composed of 36 sequences spanning 16 model organisms. These sequences were used to generate a maximum-likelihood phylogenetic tree using IQTREE version 1.6.12 (52). Branch support values were generated using ultrafast bootstrap with 1000 resamples (53). The consensus tree was selected as the final tree. The optimal substitution model for our final topology was determined to be LG (54) with invariable sites and discrete gamma model (55) based on the Bayesian Information Criterion as determined by ModelFinder (56). We rooted our final tree against an outgroup of 17 closely related human CAMK kinases using ETE Toolkit version 3.1.2 (57).

### Sequence and Structure Analysis

MAFFT (58) generated multiple sequence alignments were fed into the Bayesian Partitioning with Pattern Selection (BPPS) tool to determine evolutionarily conserved and functionally significant residues (30). Constraints mapped onto ΑlphaFold-predicted structures were visualized in PyMOL to analyze biochemical interactions.

### Rosetta Loop Modeling

Loop modeling was performed on the crystal structure (6KYQ) using the Kinematic Loop Modeling protocol (59) to model missing residues. Following this, the structure underwent five cycles of rotamer-repacking and minimization using the Rosetta Fast-relax protocol (60).

### DSSP Analysis

To analyze changes in secondary structure over our MDs, we employed the DSSP (61) command in GROMACS. This produces an output that contains an array of secondary structure values against each residue. The MDAnalysis python module (62) was used to plot these values.

### Molecular Dynamics

PDB constructs were generated by retrieving structural models RCSB and the ΑlphaFold2 database. Post-translational modifications were performed in PyMOL using the PyTMs plugin. All structures were solvated using the TIP3P water model (63). Energy minimization was run for a maximum of 10,000 steps, performed using the steepest-descent algorithm, followed by the conjugate-gradient algorithm. The system was heated from 0K to a temperature of 300K. After two equilibration steps that each lasted 20 picoseconds, 1 microsecond long simulations were run at a two femtosecond timestep. Long-range electrostatics were calculated via particle mesh Ewald (PME) algorithms using the GROMACS MD engine (64). We utilized the CHARMM36 force field (65). The resulting output was visualized using VMD 1.9.3 (66). All molecular dynamics analysis was conducted using scripts coded in Python using the MDAnalysis module (62).

### Computational Mutational Analysis

Cartesian ddG in Rosetta (67) was utilized to predict potential stabilizing and destabilizing mutations in the enzyme structure. We performed three replicates per mutation and averaged the Rosetta energies. All mutant energies were then subtracted by the wt Rosetta energy to generate a panel of ddG values relative to wt. Combined with our sequence analyses, we mutated kinase and DCLK-specific constraints to identify destabilizing interactions in the c-tail.

### Exon-Intron Boundary Mapping

The precise gene structure of DCLK1 isoforms were mapped onto the human genome with each isoform used as a query protein sequence in order to generate exon-intron borders. This was achieved using Scipio (version 1.4.1) (68) with default settings. Exons were numbered based on Ensembl annotations (69). The translation of each annotated gene sequence to protein sequence was provided with the output file (**Figure 4-source file 1-4**).

### DCLK1 cloning and recombinant protein expression

6His-DCLK1 catalytic domain (351–686), DCLK1.1 351-703 (short C-tail) or 351-729 (full C-tail), DCLK1.2 351-703 (short C-tail) or 351-740 (full C-tail), and DCLK1.2 351-703 containing D398A, N400A, V682T/V684T or D691A substitutions were synthesized by Twist Biosciences in pET28a. 6His-GST-(3C) DCLK1.1 351-689 was amplified by PCR and cloned into pOPINJ. Kinase dead, D533A 6His-GST-(3C) DCLK1.1 351-689 was generated by PCR-based site directed mutagenesis (**Figure 3-source data 3**). All plasmids were sequenced prior to their use in protein expression studies. All proteins, including 6His-GST-(3C) DCLK1 351-689, with a 3C-protease cleavable affinity tag, were expressed in BL21(DE3)pLysS *E. coli* (Novagen) and purified by affinity and size exclusion chromatography. The short N-terminal 6-His affinity tag present on all other DCLK1 proteins described in this paper was left in situ on recombinant proteins, since it does not appear to interfere with DSF, biochemical interactions or catalysis. For analytical SEC chromatography, 1 mg of each DCLK1 protein was assayed on a Superdex 200 Increase 10/300 GL (Cytiva), and the eluted fractions were also analysed by SDS-PAGE and Coomassie blue staining to confirm composition. The molecular weight standards were loaded in a mixture of 200 ug of Bovine Serum Albumin (BSA), Carbonic Anhydrase (CA), and Alcohol Dehydrogenase (AD) each.

### Mass Spectrometry

Purified DCLK1 proteins (5 μg) were diluted (∼40-fold) in 100 mM ammonium bicarbonate pH 8.0 and reduced (DTT) and alkylated (iodoacetamide, as previously described (70), and digested with a 25:1 (w/w) trypsin gold (Promega) at 37 °C for 18 hours with gentle agitation. Digests were then subjected to strong cation exchange chromatography using in-house packed stage tip clean-up (71). Dried tryptic peptides were solubilized in 20 μl of 3% (v/v) acetonitrile and 0.1% (v/v) TFA in water, sonicated for 10 min, and centrifuged at 13,000 x *g* for 10 min at 4°C and supernatant collected. LC-MS/MS separation was performed using an Ultimate 3000 nano system (Dionex), over a 60-min gradient (70). Briefly, samples were loaded at a rate of 12 μL/min onto a trapping column (PepMap100, C18, 300 μm × 5 mm) in loading buffer (3% (v/v) acetonitrile, 0.1% (v/v) TFA). Samples were then resolved on an analytical column (Easy-Spray C18 75 μm × 500 mm, 2 μm bead diameter column) using a gradient of 97% A (0.1% (v/v) formic acid): 3% B (80% (v/v) acetonitrile, 0.1% (v/v) formic acid) to 80% B over 30 min at a flow rate of 300 nL/min. All data acquisition was performed using a Fusion Lumos Tribrid mass spectrometer (Thermo Scientific). Samples were injected twice with either higher-energy C-trap dissociation (HCD) fragmentation (set at 32% normalized collision energy [NCE]) or Electron transfer dissociation (ETD) with supplemental 30% NCE HCD (EThcD) for 2+ to 4+ charge states using a top 3s top speed mode. MS1 spectra were acquired at a 120K resolution (at 200 *m/z*), over a range of 300 to 2000 *m/z*, normalised AGC target = 50%, maximum injection time = 50 ms. MS2 spectra were acquired at a 30K resolution (at 200 *m/z*), AGC target = standard, maximum injection time = dynamic. A dynamic exclusion window of 20 s was applied at a 10 ppm mass tolerance. Data was analysed by Proteome Discoverer 2.4 in conjunction with the MASCOT search engine using a custom database of the UniProt *Escherichia coli* reviewed database (Updated January 2023) with the DCLK1 mutant variant amino acid sequences manually added, and using the search parameters: fixed modification = carbamidomethylation (C), variable modifications = oxidation (M) and phospho (S/T/Y), MS1 mass tolerance = 10 ppm, MS2 mass tolerance = 0.01 Da, and the *ptm*RS node on; set to a score > 99.0. For HCD data, instrument type = electrospray ionization–Fourier-transform ion cyclotron resonance (ESI-FTICR), for EThcD data, instrument type = EThcD. For label free relative quantification of phosphopeptide abundances of the different DCLK1 variants, the minora feature detector was active and set to calculate the area under the curve for peptide *m/z* ions. Abundance of phosphopeptide ions were normalised against the total protein abundance (determine by the HI3 method (72), as in the minora feature detector node) to account for potential protein load variability during analysis.

### DCLK1 DSF

Thermal Shift Assays (TSA), were performed using Differential Scanning Fluorimetry (DSF) in a StepOnePlus Real-Time PCR machine (Life Technologies) in combination with Sypro-Orange dye (Invitrogen) and a thermal ramping protocol (0.3°C per minute between 25 and 94°C). Recombinant DCLK1 proteins were assayed at a final concentration of 5 μM in 50 mM Tris–HCl (pH 7.4) and 100 mM NaCl in the presence or absence of the indicated concentrations of ligand (ATP or Mg:ATP) or DCLK1 inhibitor compounds, with final DMSO concentrations never higher than 4% (v/v). Thermal melting data were processed using the Boltzmann equation to generate sigmoidal denaturation curves, and average *T*_m_/Δ*T*_m_ values were calculated as described using GraphPad Prism software, as previously described, from 3 technical repeats (73).

### DCLK1 kinase assays

DCLK1 peptide-based enzyme assays (74,75) were carried out using the LabChip EZ Reader platform, which monitors and quantifies real-time phosphorylation-induced changes in the mobility of the fluorescently-labelled DCLK1 peptide substrate 5-FAM-KKALRRQETVDAL-CONH_2_. To assess DCLK1 catalytic domains, or DCLK1.1 or DCLK1.2 variants, 100ng of purified protein were incubated with a high (1 mM) concentration of ATP (to mimic cellular levels of nucleotide) and 2 μM of the fluorescent substrate in 25 mM HEPES (pH 7.4), 5 mM MgCl_2_, and 0.001% (v/v) Brij 35. DCLK1-IN-1 and DCLK1-NEG (kind gifts from Dr Fleur Ferguson, UCSF) enzyme inhibition was quantified under identical assay conditions in the presence of 10 μM of each compound. Assays are either reported as rates (pmoles/min phosphate incorporation) during linear phosphate incorporation (e.g total substrate phosphorylation limited to <20-30% to prevent ATP depletion and to ensure assay linearity), or presented as time-dependent percentage substrate phosphorylation (kinetic mode). Rates of substrate phosphorylation (pmol phosphate incorporation per min) were determined using a fixed amount of kinase and linear regression analysis with GraphPad Prism software; V_max_ and *K*_M[ATP]_ values were calculated at 2 µM substrate peptide concentration, as previously described (76). Rates are normalized to enzyme concentration and all enzyme rate and kinetic data are presented as mean and SD of 4 technical replicates.

### Author Contributions

N.K. conceptualization; A.V., G.W., D.P.B., C.E.E., P.A.E., and N.K. methodology; A.V., G.W., D.P.B., B.O., S.S., N.G., E.E.F., and L.A.D. investigation; A.V., G.W., and B.O., W.Y. data curation; A.V., G.W., D.P.B., B.O., and S.S. formal analysis; A.V., G.W., D.P.B.., and N.G. validation; A.V., G.W., D.P.B., P.A.E., and N.K. writing – original draft; A.V., G.W., D.P.B., B.O., S.S., N.G., E.E.F., L.A.D., C.B., W.Y., I.A., S.K., C.E.E., P.A.E., and N.K. writing – review and editing; A.V., G.W., D.P.B., B.O., and S.S. visualization; P.A.E. and N.K. supervision; A.V., D.P.B., E.E.F., L.A.D., C.E.E., P.A.E., and N.K. funding acquisition.

### Funding and additional information

Funding from N.K. (grant no.: R35 GM139656) is acknowledged. P.A.E. acknowledges funding from a University of Liverpool BBSRC IAA award. A.V. acknowledges funding from ARCS Foundation. D.P.B, L.A.D., P.A.E. and C.E.E. also acknowledge BBSRC grants BB/S018514/1, BB/N021703/1 and BB/X002780/1 and North West Cancer Research (NWCR) grant CR1208. E.E.F. thanks the MRC for a DiMeN DTP studentship (No. 1961582). The content is solely the responsibility of the authors and does not necessarily represent the official views of the National Institutes of Health.

### Data Availability

All data generated in this study are included within the manuscript. Source data are provided for each figure. MD simulations and associated data may be accessed from https://www.dropbox.com/sh/xtiwpjgyzxy1oz0/AACK6dS3ypzYXDih3wgKp9bla?dl=0.

## Figure supplements for

**Figure 1-figure supplement 1:**
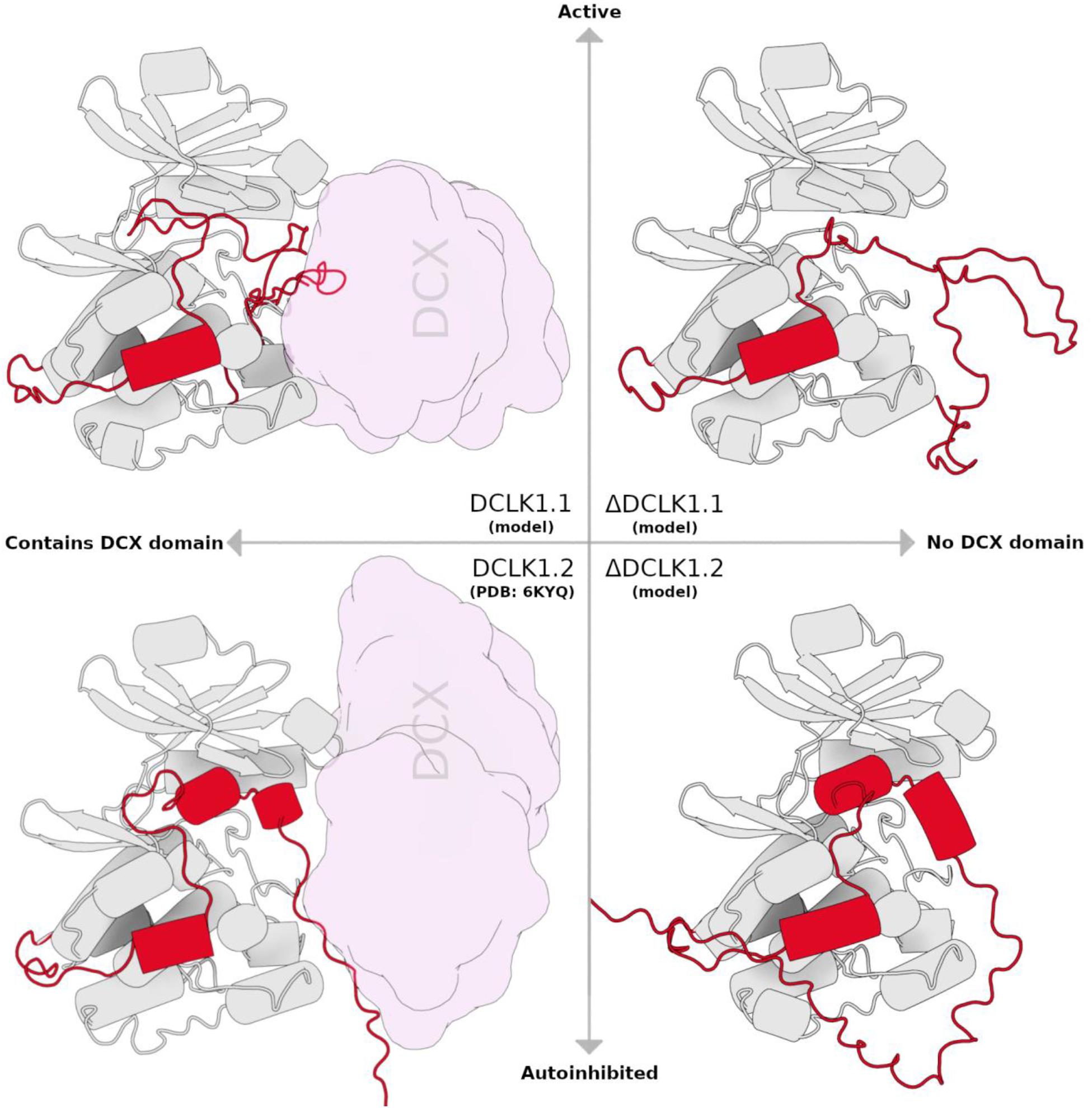
Structural cartoon depicting each DCLK1 isoform, categorized by the presence of DCX domain and related to enzymatic activity

**Figure 2-figure supplement 1:**
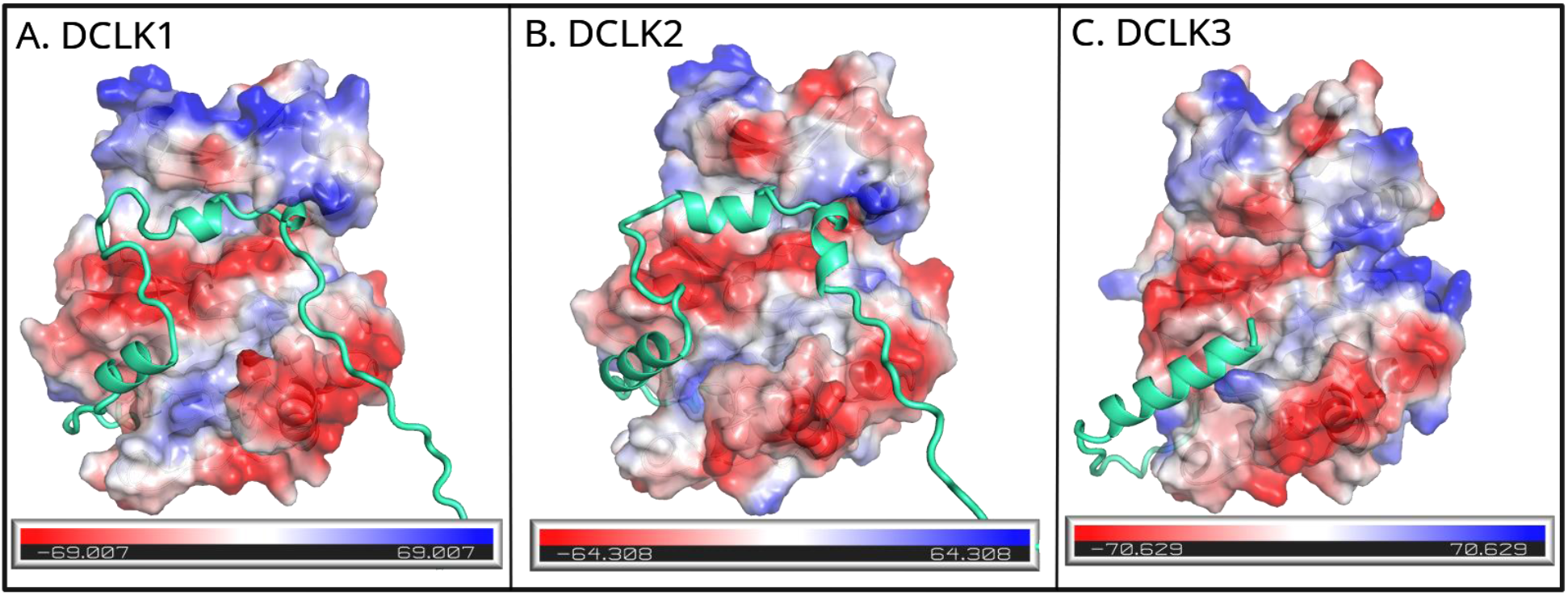
Electrostatic surface views of full-length DCLK paralog (DCLK1, DCLK2, and DCLK3) in the same orientation. These structures show how the tail packs against the substrate binding pocket of the kinase domain in each paralog. The electrostatic surface is color-coded with negative (red) and positive (blue) charges. Similarities in the distribution of charges among paralogs can influence substrate binding affinity and kinase activity.

**Figure 3-figure supplement 1:**
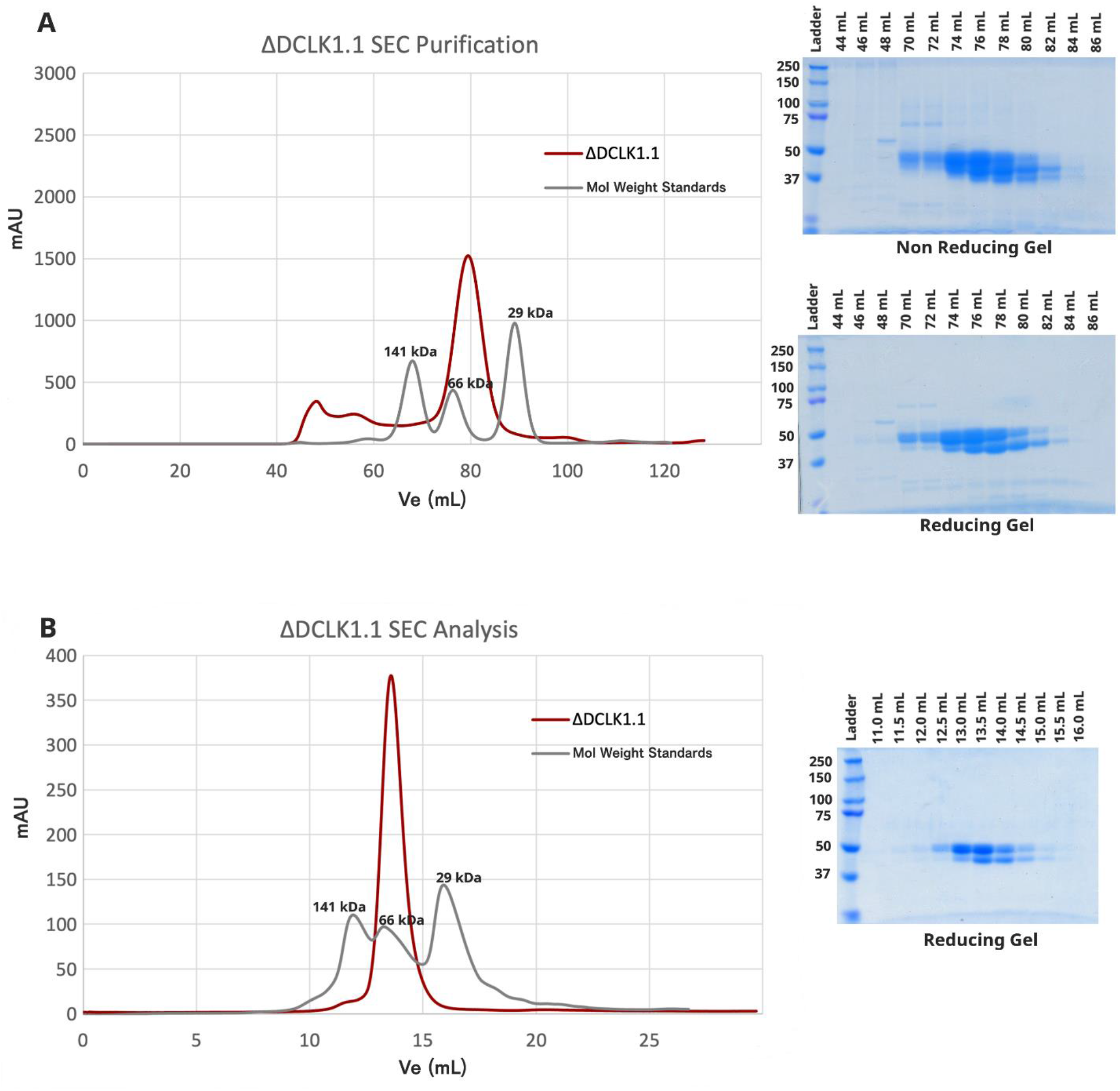
**A**) Purification of DCLK1.1^351-729^ by SEC reveals a single peak, representing monomeric protein in solution. Reducing and non-reducing SDS-PAGE show the elution profiles of the purified species. **B)** Analytical SEC of purified DCLK1.1^351-729^ (1.1 mg) confirms a single monomeric species.

**Figure 3-figure supplement 2:**
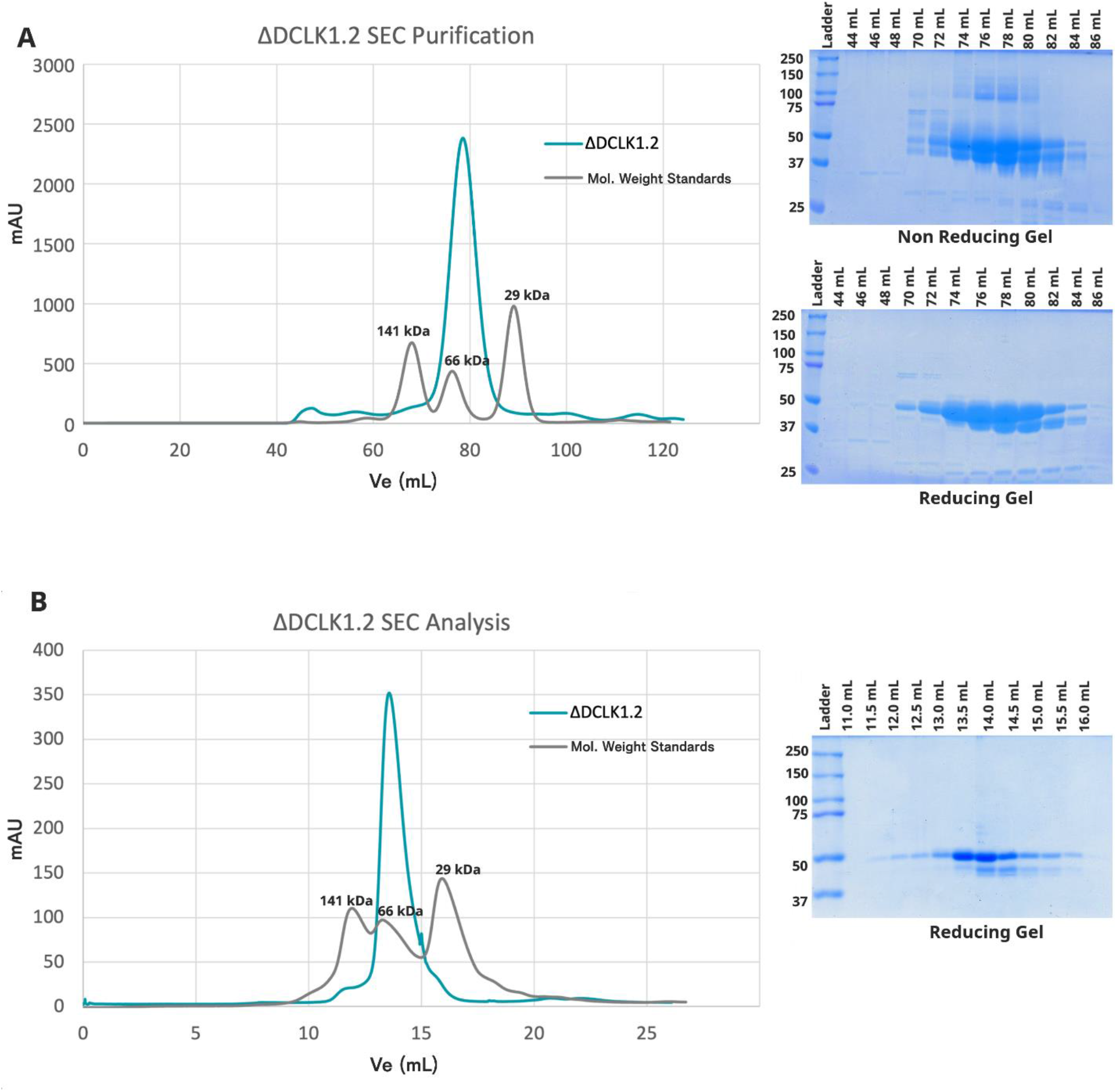
**A**) Purification of DCLK1.2^351-740^ by SEC reveals a single peak, representing monomeric protein in solution. Reducing and non-reducing SDS-PAGE showing the elution profiles of the purified proteins. **B)** Analytical SEC of purified DCLK1.2^351-740^ (1.1 mg) confirms a single monomeric species.

**Figure 3-figure supplement 3:**
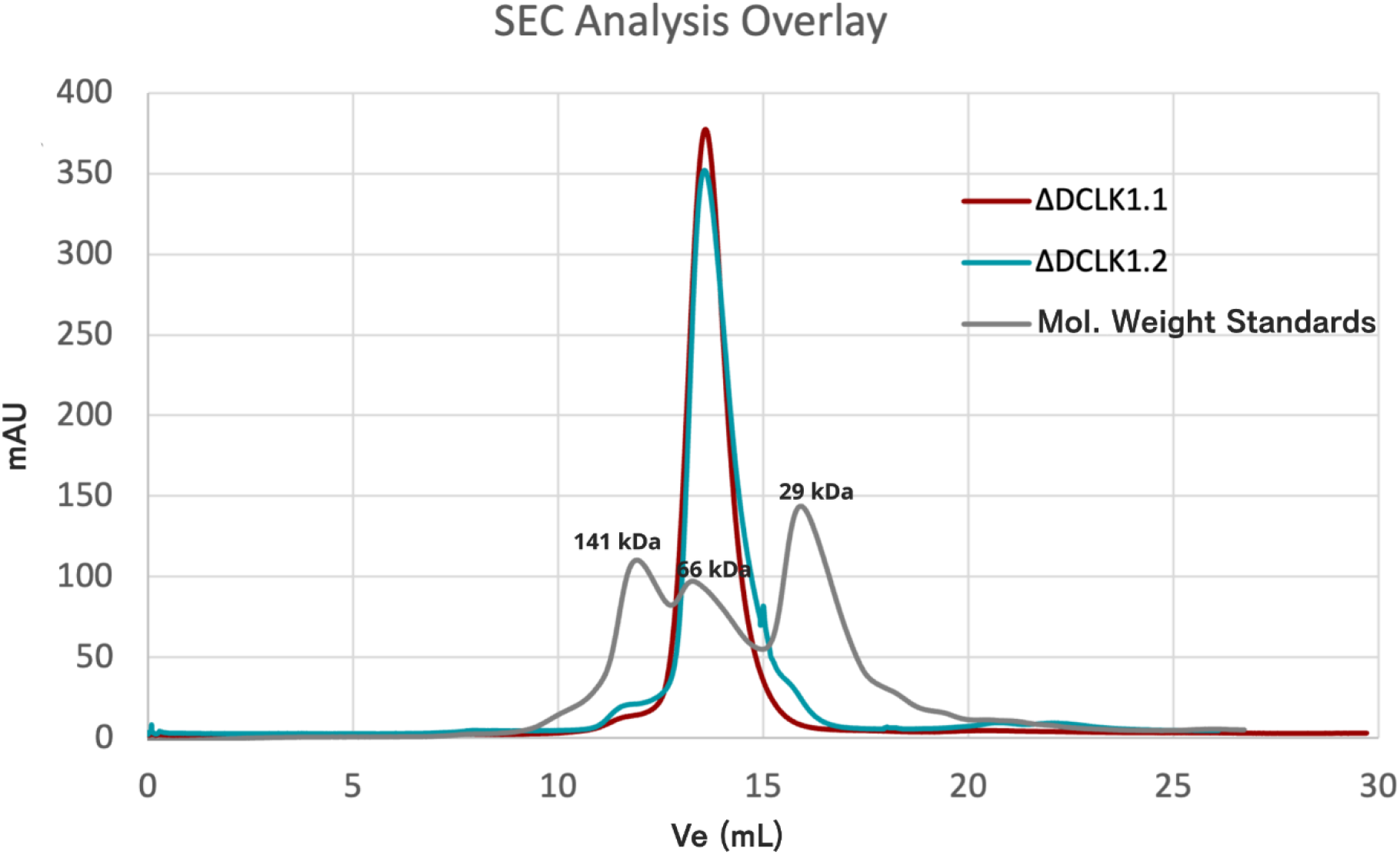
Analytical SEC of DCLK1.1^351-729^ and DCLK1.2^351-740^. Overlaid chromatograms showing elution profiles for both proteins and molecular weight standards. Both DCLK1 isoforms elute as single peaks at the predicted molecular weight for the monomeric protein.

**Figure 3-figure supplement 4:**
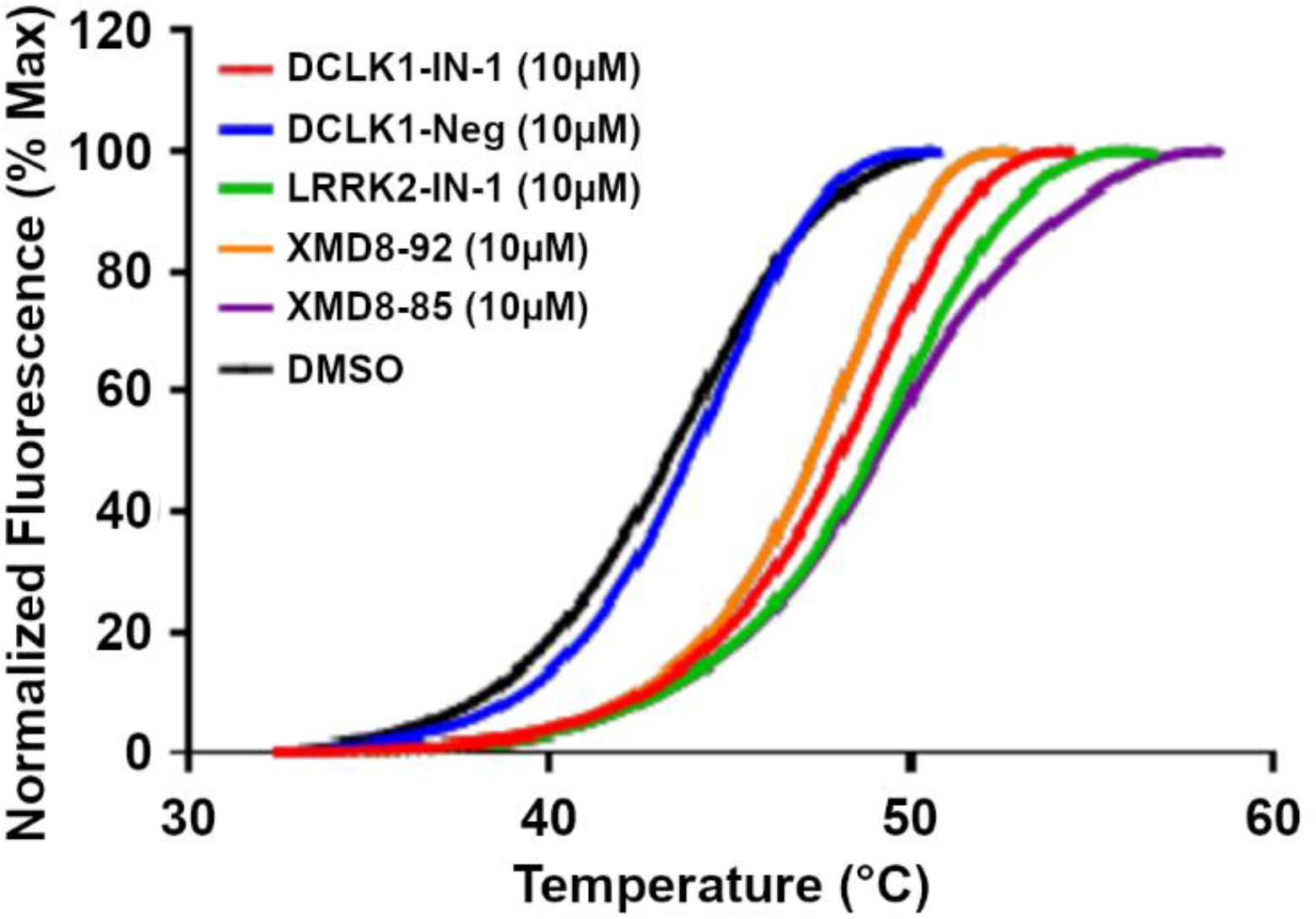
DSF profile of DCLK1_351-689_ in the presence of DMSO or a panel of DCLK1 inhibitor compounds. DCLK1-Neg is a negative control.

**Figure 5-figure supplement 1:**
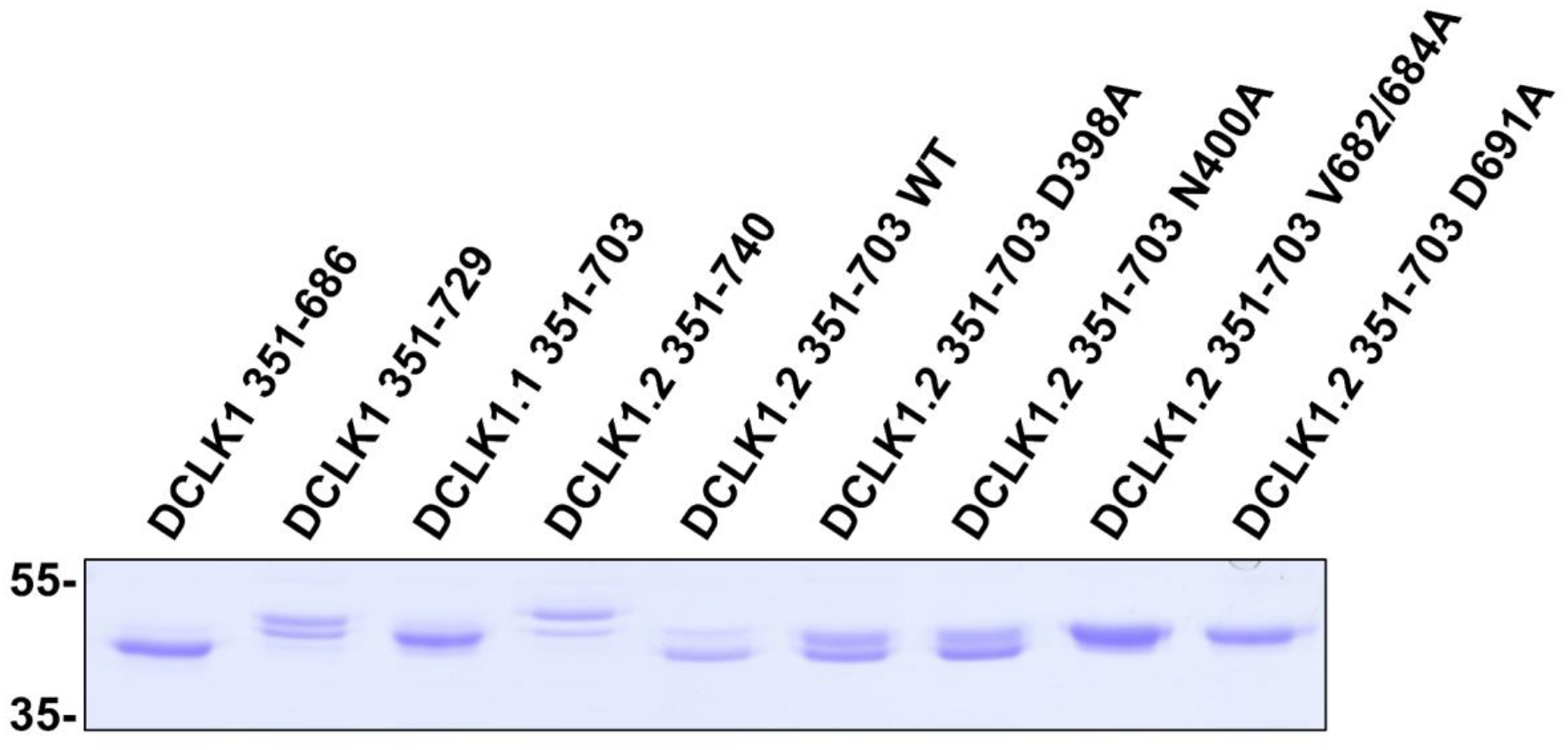
SDS-Page and Coomassie blue staining of each DCLK1 protein.

**Figure 5-figure supplement 2:**
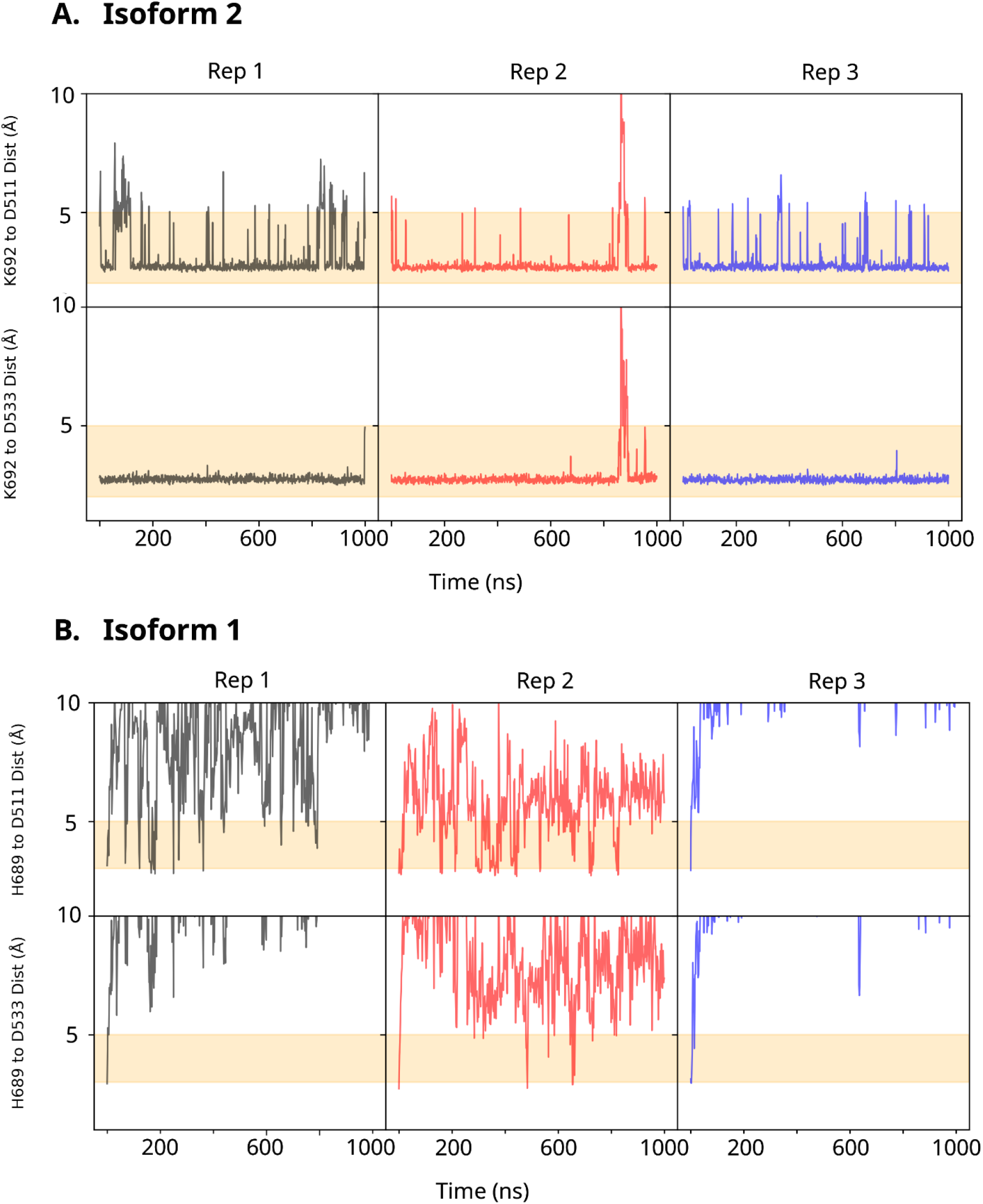
**A**) Minimum Distance of K692 in the DCLK1.2 C-tail forms significant stable interactions over microsecond replicates to the DFG and HRD aspartates. **B)** H689 in the DCLK1.1 C-tail, comparatively fails to interact with the DFG and HRD aspartates.

**Figure 5-figure supplement 3:**
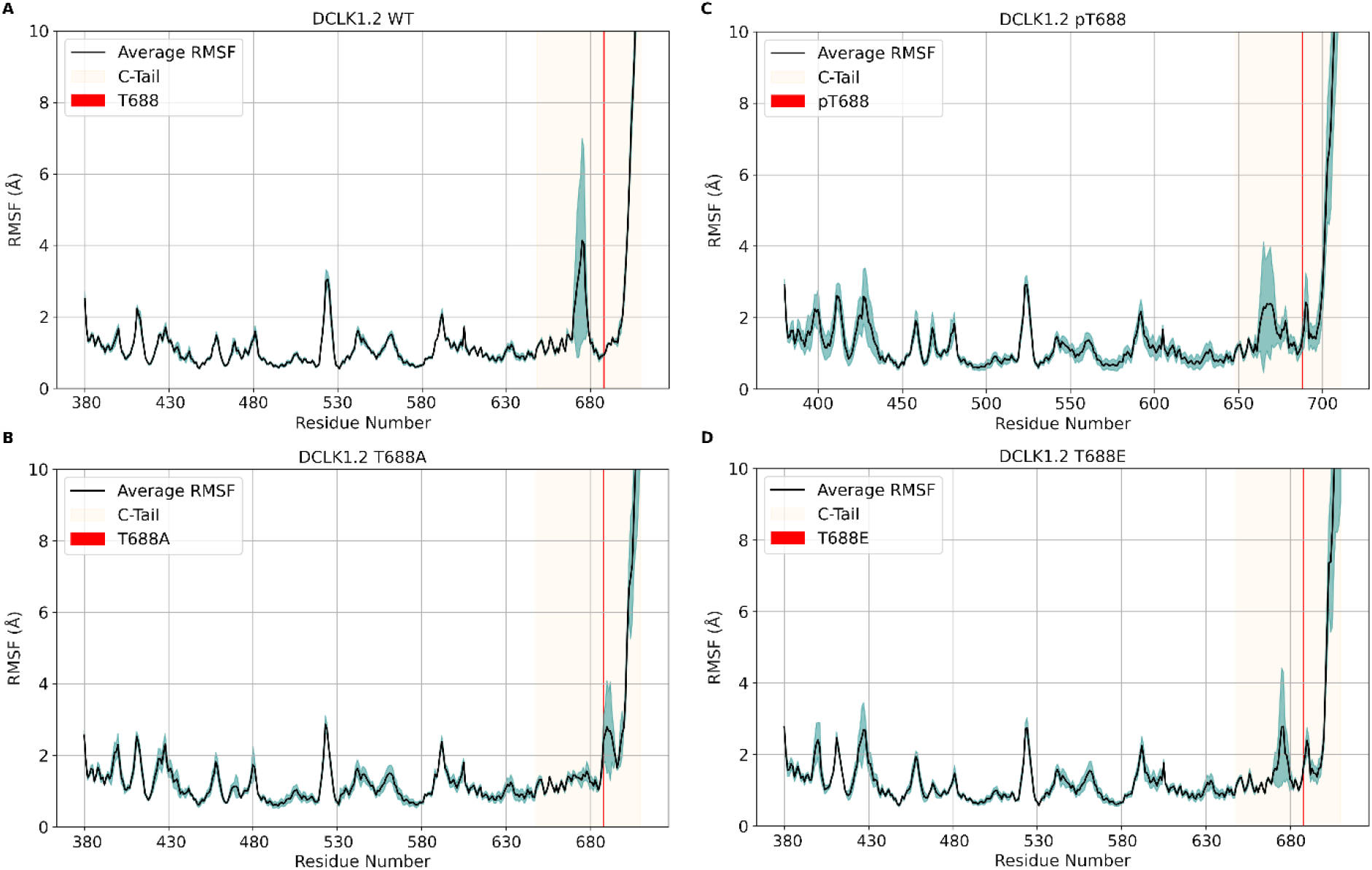
RMSF plots of MD simulations of ΔDCLK1.2 wt, pT688, T688A, and T688E, where T688 is demarcated by a red line and the entire ΔDCLK1.2 C-tail is highlighted in light yellow. The black line represents the average RMSF between three 500ns replicates and the blue shading represents standard deviation of the replicates, where less shading indicates convergence between replicates and increased shading indicates deviation.

**Figure 6-figure supplement 1:**
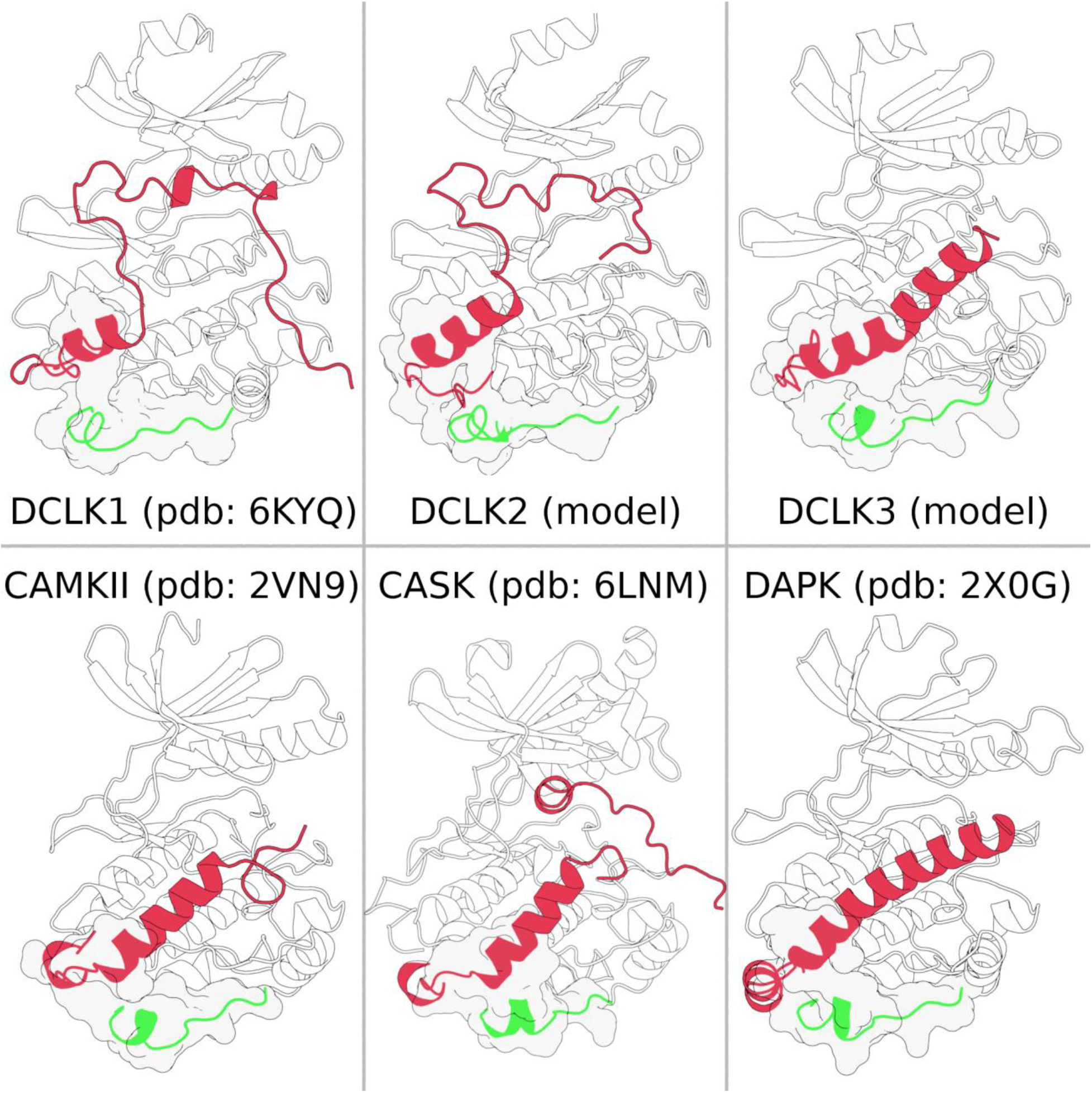
CAMK-specific insert (green) consistently making structural contacts (shown in surface representation) with the C-tail (red) across multiple CAMK families.

**Figure 6-figure supplement 2:**
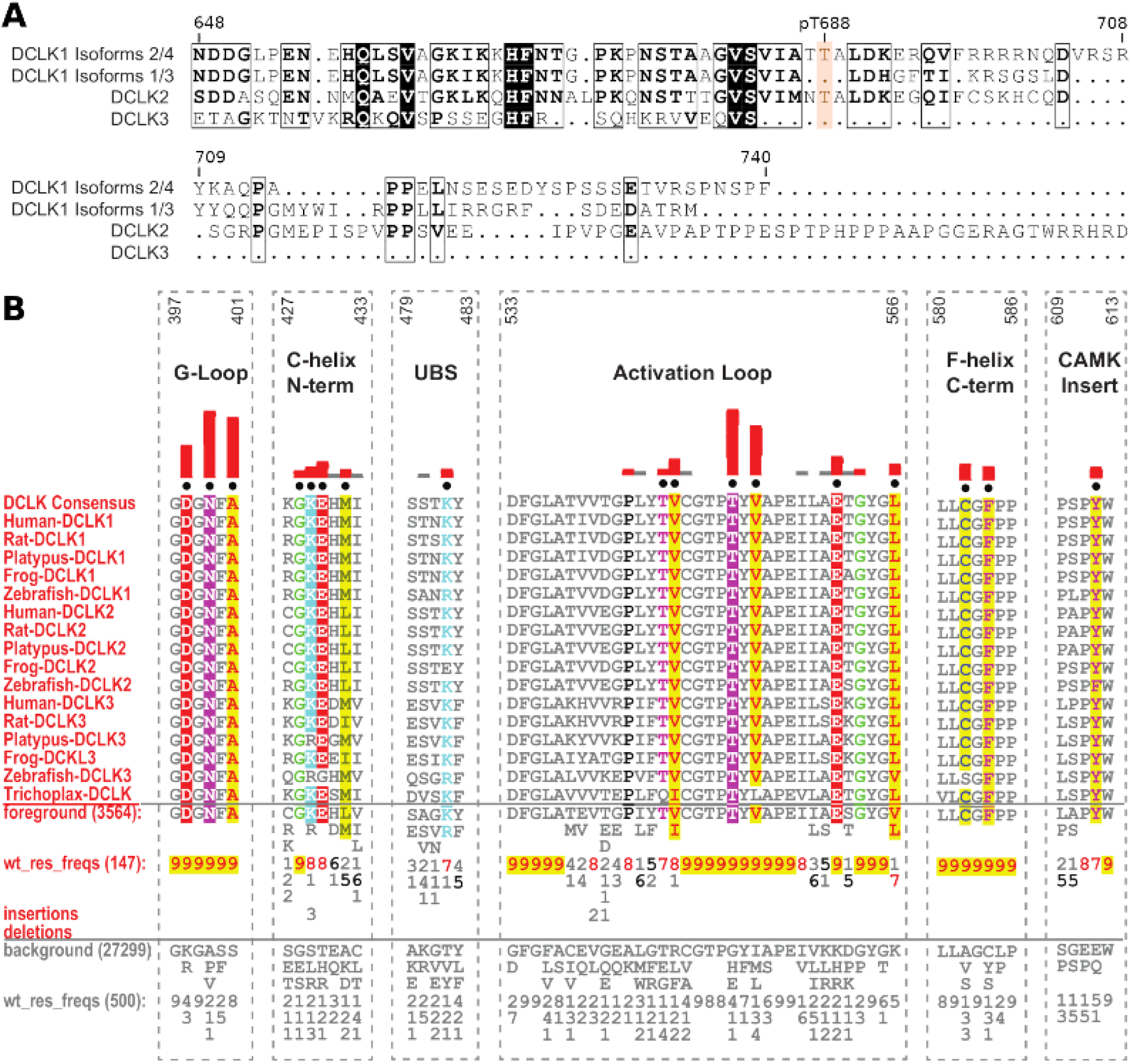
Identification of DCLK family-specific constraints. **A**) Sequence alignment of human DCLK paralogs including long and short isoforms of DCLK1. **B)** Sequence constraints that distinguish DCLK1/2/3 sequences from closely related CAMK sequences are shown in a contrast hierarchical alignment (CHA). The CHA shows DCLK1/2/3 sequences from diverse organisms as the display alignment. The foreground consists of 3564 DCLK sequences while the background alignment contains 27,299 related CAMK sequences. The foreground and background alignments are shown as residue frequencies below the display alignment in integer tenths (1–9). The histogram (red) indicates the extent to which distinguishing residues in the foreground diverge from the corresponding position in the background alignment. Black dots indicate the alignment positions used by the BPPS (Neuwald, 2014) procedure when classifying DCLK sequences from related CAMK sequences. Alignment number is based on the human DCLK1 sequence (Uniprot ID: O15075-1).

**Figure 7-figure supplement 1:**
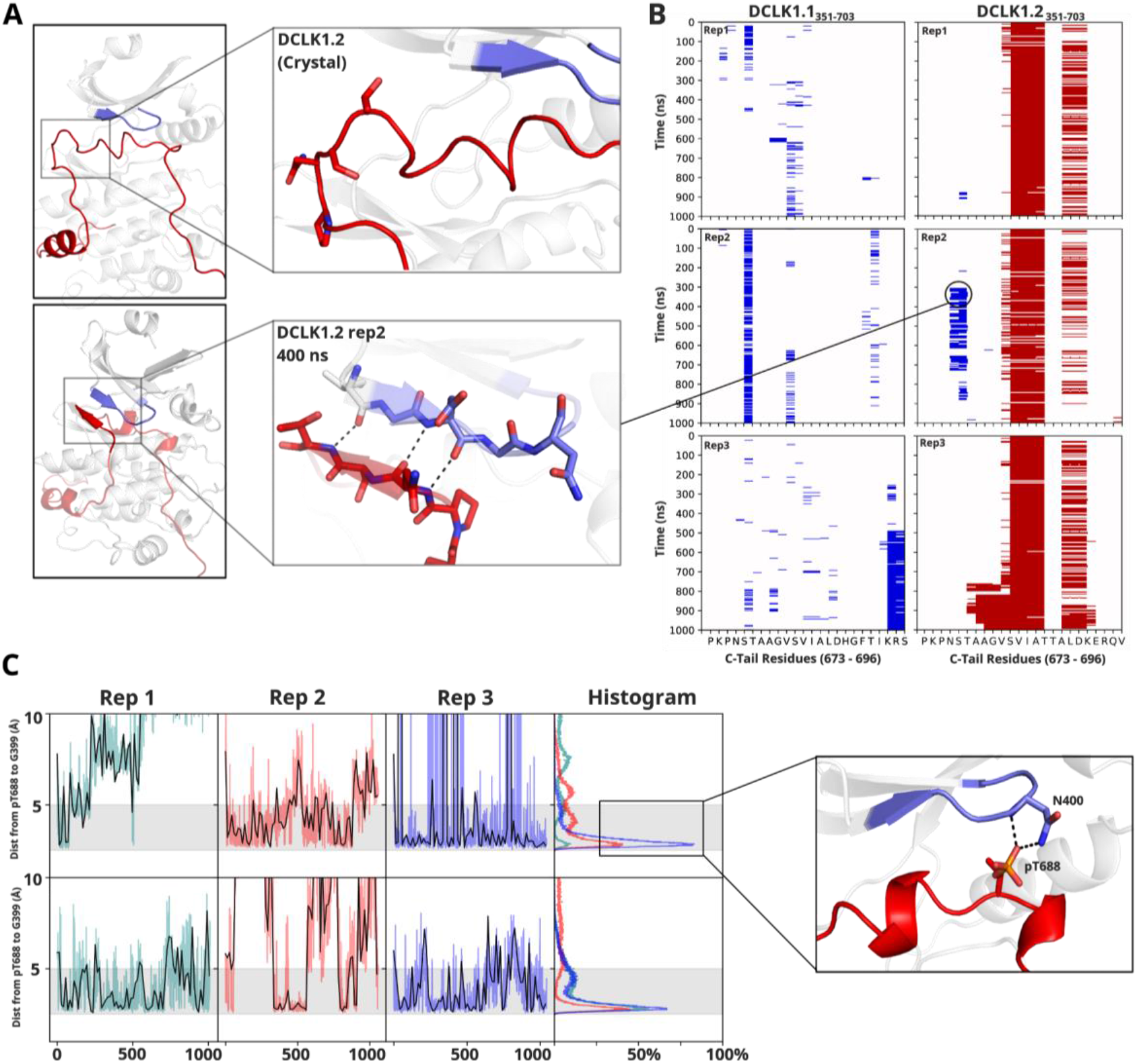
Molecular Dynamics of DCLK1 isoforms. **A-B)** Microsecond MD replicates from DCLK1.1 and DCLK1.2, showing the DSSP output plotted for the C-tail, where red lines represent alpha helices and blue lines represent beta sheets. **C)** Distance plots from MD replicates of the phosphorylated threonine highlighting the contact distance between pT688 phosphate and G399 of the G-loop.

**Figure 8-figure supplement 1:**
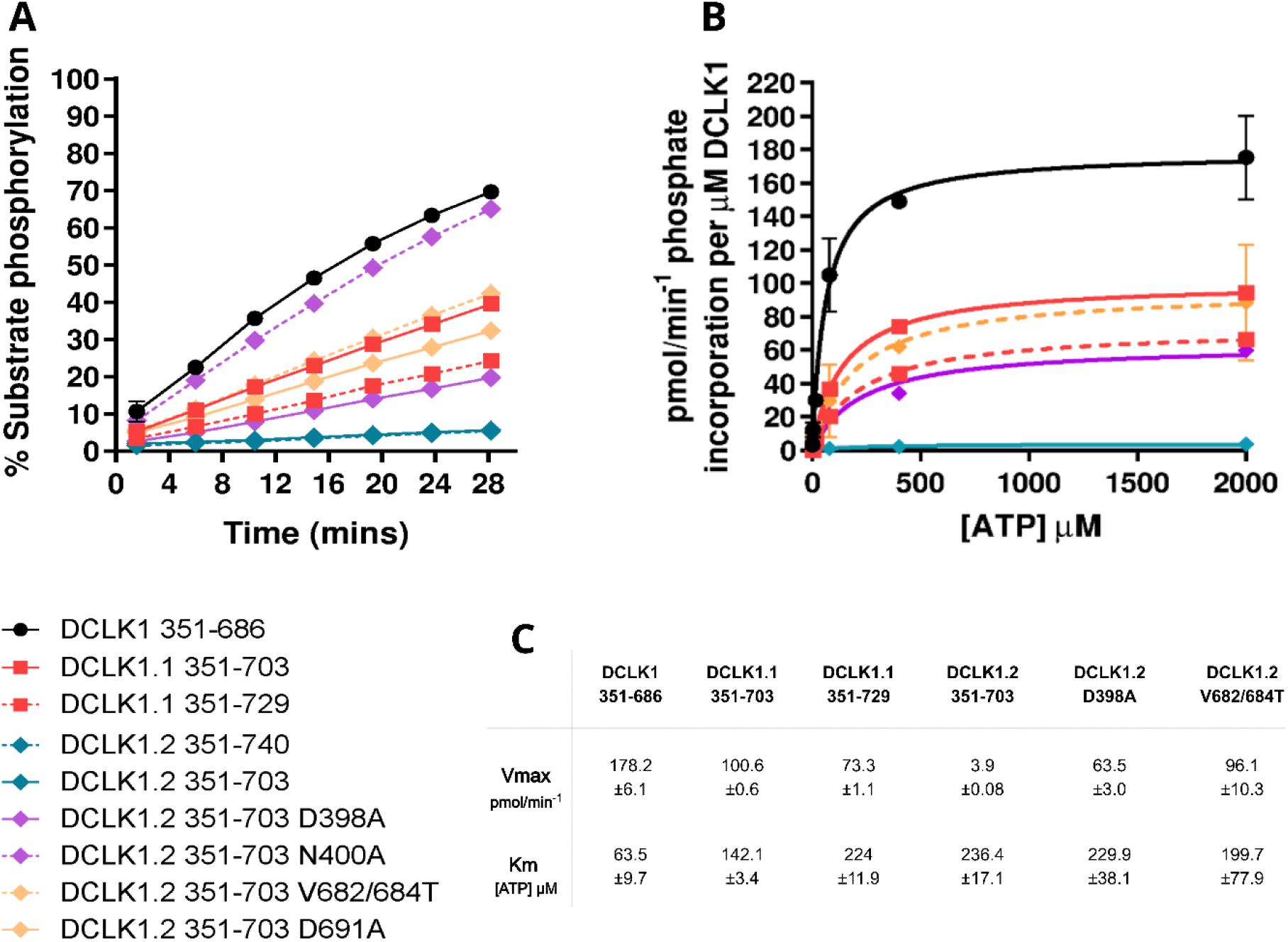
**A**) DCLK1 substrate phosphorylation (calculated as % total phosphopeptide) was quantified as a function of time for each of the indicated purified DCLK proteins in the presence of 1 mM ATP. Assays were performed side-by-side. Data is mean and SD from (N=4) independent experiments. **B)** Michaelis-Menten plots showing normalized DCLK1 activity in the presence of increasing concentrations of ATP, to tease apart effects of C-tail on ATP affinity. Data shown is mean and SD from (N=3) independent experiments. **C)** Table of calculated Vmax and Km [ATP] values obtained from (B).

**Figure 8-figure supplement 2:**
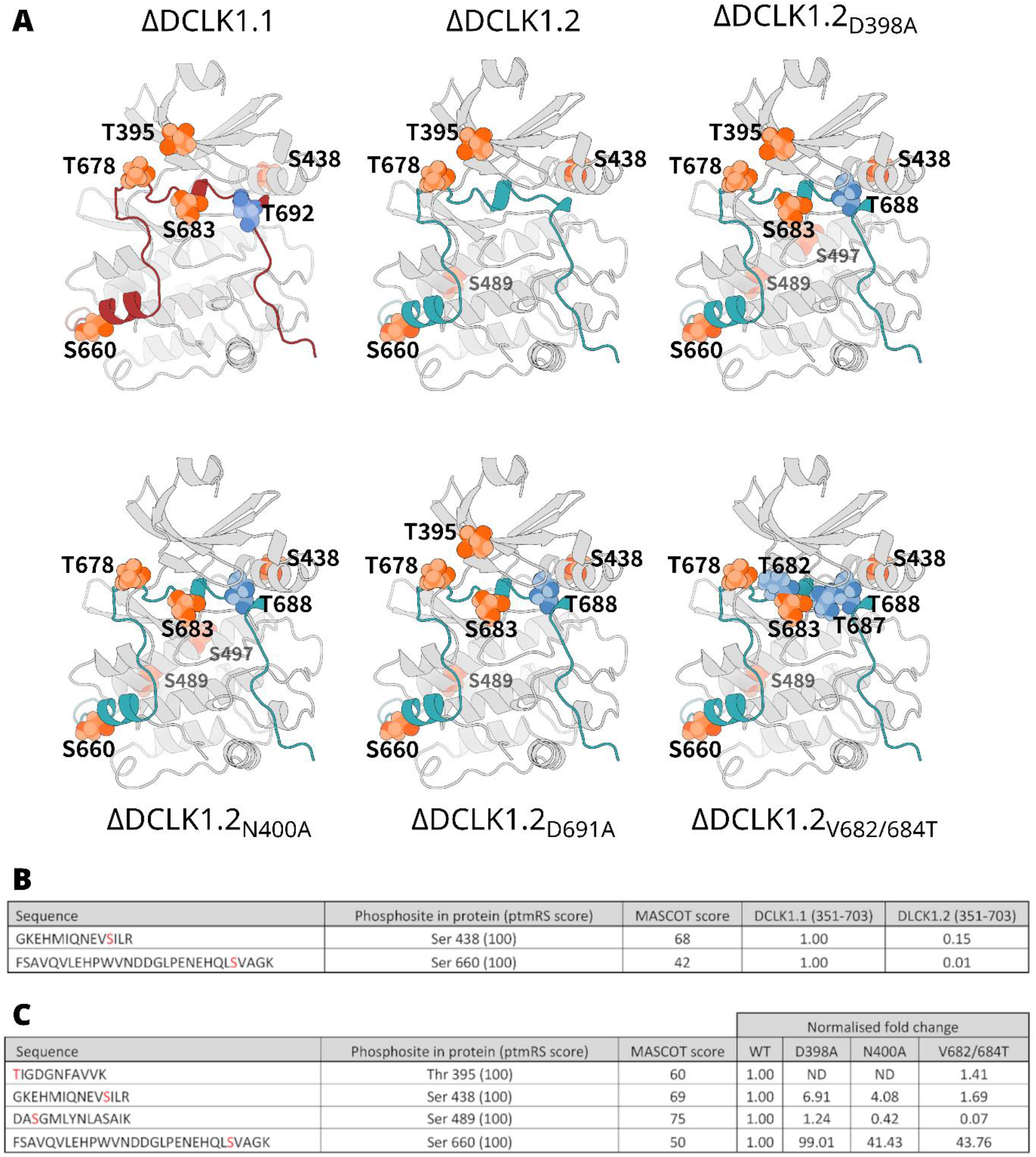
**A**) All mapped DCLK phosphorylation sites derived from LC-MS/MS analysis of DCLK1.1 and DCLK 1.2 proteins. Identified sites of phosphorylation at the kinase domain are colored in orange with isoform or mutant-specific phosphorylation sites colored in blue and mapped onto the structure of each protein **B)** Quantitative LC-MS/MS data showing tryptic phosphopeptides identified from DCLK1.1 and 1.2 that were directly comparable between isoforms. Detailed are peptide sequences, identified sites of phosphorylation (red), the site of phosphorylation within the protein polypeptide and the ptmRS score relevant to confidence of phosphosite localisation, as well as the Mascot score for peptide identification. Fold-changes in the relative abundance of the two phosphopeptides in DCLK 1.2 are computed with reference to these same two phosphopeptides in DCLK1.1, normalising against 3 non-modified peptides to account for potential difference in the amount analysed. **C)** As described in B, quantitative LC-MS/MS data for sites directly comparable between DCLK1.2 and its variants. Fold change in abundance could not be calculated for the peptide containing pThr 395 given the presence of the inserted amino acid mutations and the differences in relative ionisation efficiency for the resulting tryptic peptide.

**Figure 9-figure supplement 1:**
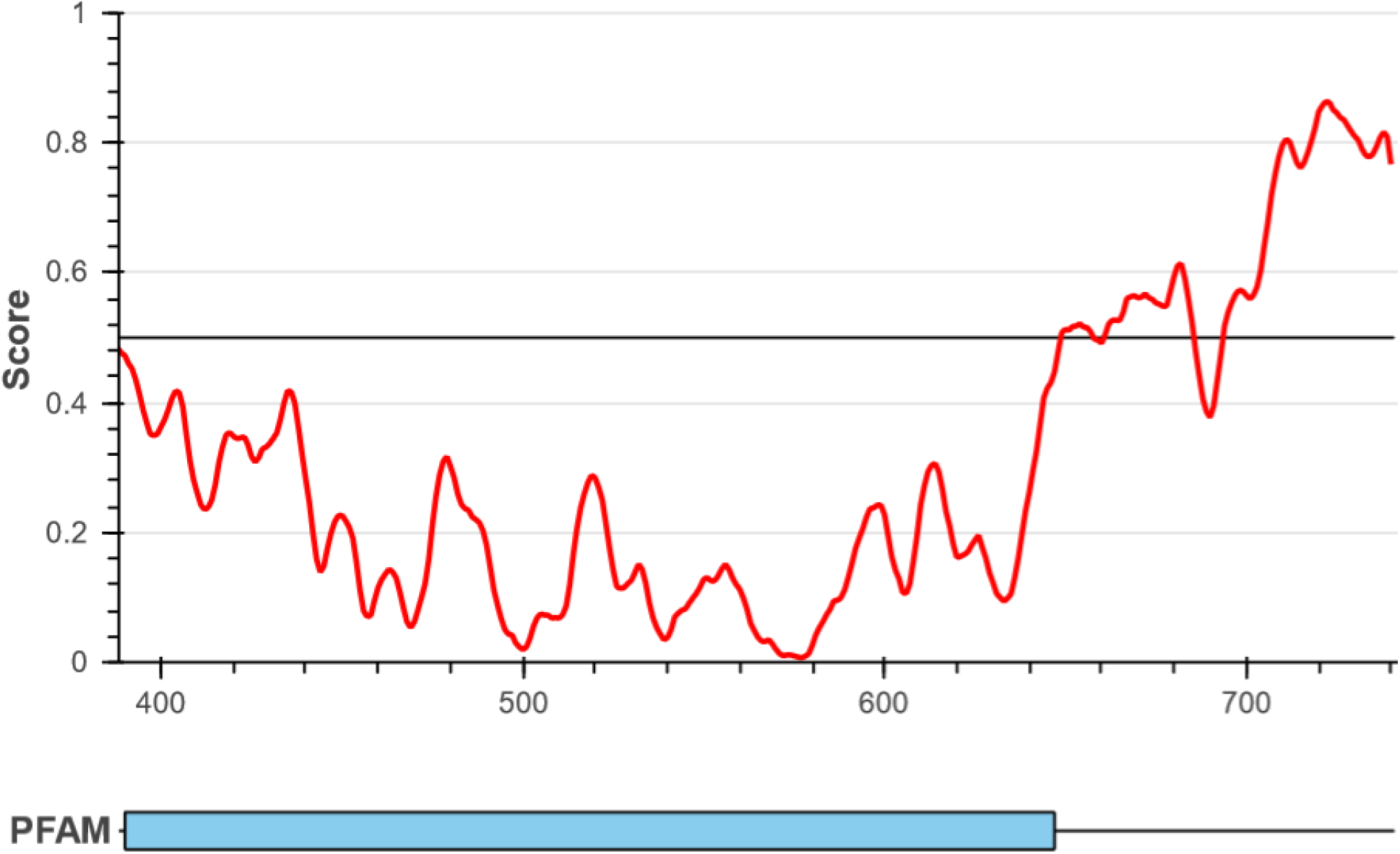
Intrinsic Disorder prediction of DCLK1.2 C-tail using IUPRED3.

## References

1. Manning G, Whyte DB, Martinez R, Hunter T, Sudarsanam S. The protein kinase complement of the human genome. Science. 2002 Dec 6;298(5600):1912–34.

2. Agulto RL, Rogers MM, Tan TC, Ramkumar A, Downing AM, Bodin H, et al. Autoregulatory control of microtubule binding in doublecortin-like kinase 1. Carter AP, Sengupta P, editors. eLife. 2021 Jul 26;10:e60126.

3. Bayer KU, Schulman H. CaM Kinase: Still Inspiring at 40. Neuron. 2019 Aug 7;103(3):380–94.

4. Gógl G, Kornev AP, Reményi A, Taylor SS. Disordered Protein Kinase Regions in Regulation of Kinase Domain Cores. Trends Biochem Sci. 2019 Apr;44(4):300–11.

5. Berginski ME, Moret N, Liu C, Goldfarb D, Sorger PK, Gomez SM. The Dark Kinase Knowledgebase: an online compendium of knowledge and experimental results of understudied kinases. Nucleic Acids Res. 2021 Jan 8;49(D1):D529–35.

6. Sossey-Alaoui K, Srivastava AK. DCAMKL1, a brain-specific transmembrane protein on 13q12.3 that is similar to doublecortin (DCX). Genomics. 1999 Feb 15;56(1):121–6.

7. Ohmae S, Takemoto-Kimura S, Okamura M, Adachi-Morishima A, Nonaka M, Fuse T, et al. Molecular identification and characterization of a family of kinases with homology to Ca2+/calmodulin-dependent protein kinases I/IV. J Biol Chem. 2006 Jul 21;281(29):20427–39.

8. Couillard-Despres S, Winner B, Schaubeck S, Aigner R, Vroemen M, Weidner N, et al. Doublecortin expression levels in adult brain reflect neurogenesis. European Journal of Neuroscience. 2005;21(1):1–14.

9. Horesh D, Sapir T, Francis F, Grayer Wolf S, Caspi M, Elbaum M, et al. Doublecortin, a Stabilizer of Microtubules. Human Molecular Genetics. 1999 Sep 1;8(9):1599–610.

10. Cheng L, Huang S, Chen L, Dong X, Zhang L, Wu C, et al. Research Progress of DCLK1 Inhibitors as Cancer Therapeutics. Curr Med Chem. 2022;29(13):2261–73.

11. Gao T, Wang M, Xu L, Wen T, Liu J, An G. DCLK1 is up-regulated and associated with metastasis and prognosis in colorectal cancer. J Cancer Res Clin Oncol. 2016 Oct 1;142(10):2131–40.

12. Westphalen CB, Quante M, Wang TC. Functional implication of Dclk1 and Dclk1-expressing cells in cancer. Small GTPases. 2017 Jul 3;8(3):164–71.

13. Galvan L, Francelle L, Gaillard MC, de Longprez L, Carrillo-de Sauvage MA, Liot G, et al. The striatal kinase DCLK3 produces neuroprotection against mutant huntingtin. Brain. 2018 May 1;141(5):1434– 54.

14. Kannan N, Haste N, Taylor SS, Neuwald AF. The hallmark of AGC kinase functional divergence is its C-terminal tail, a cis-acting regulatory module. Proc Natl Acad Sci U S A. 2007 Jan 23;104(4):1272– 7.

15. Yeon JH, Heinkel F, Sung M, Na D, Gsponer J. Systems-wide Identification of cis-Regulatory Elements in Proteins. Cell Syst. 2016 Feb 24;2(2):89–100.

16. Nguyen T, Ruan Z, Oruganty K, Kannan N. Co-conserved MAPK features couple D-domain docking groove to distal allosteric sites via the C-terminal flanking tail. PLoS One. 2015;10(3):e0119636.

17. Yeung W, Kwon A, Taujale R, Bunn C, Venkat A, Kannan N. Evolution of Functional Diversity in the Holozoan Tyrosine Kinome. Mol Biol Evol. 2021 Dec 9;38(12):5625–39.

18. Kwon A, John M, Ruan Z, Kannan N. Coupled regulation by the juxtamembrane and sterile α motif (SAM) linker is a hallmark of ephrin tyrosine kinase evolution. J Biol Chem. 2018 Apr 6;293(14):5102–16.

19. Rellos P, Pike ACW, Niesen FH, Salah E, Lee WH, von Delft F, et al. Structure of the CaMKIIdelta/calmodulin complex reveals the molecular mechanism of CaMKII kinase activation. PLoS Biol. 2010 Jul 27;8(7):e1000426.

20. Bhattacharyya M, Lee YK, Muratcioglu S, Qiu B, Nyayapati P, Schulman H, et al. Flexible linkers in CaMKII control the balance between activating and inhibitory autophosphorylation. Elife. 2020 Mar 9;9:e53670.

21. Hudmon A, Schulman H. Structure-function of the multifunctional Ca2+/calmodulin-dependent protein kinase II. Biochem J. 2002 Jun 15;364(Pt 3):593–611.

22. Huse M, Kuriyan J. The conformational plasticity of protein kinases. Cell. 2002 May 3;109(3):275–82.

23. Wayman GA, Lee YS, Tokumitsu H, Silva AJ, Soderling TR. Calmodulin-kinases: modulators of neuronal development and plasticity. Neuron. 2008 Sep 25;59(6):914–31.

24. Omori Y, Suzuki M, Ozaki K, Harada Y, Nakamura Y, Takahashi E, et al. Expression and chromosomal localization of KIAA0369, a putative kinase structurally related to Doublecortin. J Hum Genet. 1998;43(3):169–77.

25. Matsumoto N, Pilz DT, Ledbetter DH. Genomic structure, chromosomal mapping, and expression pattern of human DCAMKL1 (KIAA0369), a homologue of DCX (XLIS). Genomics. 1999 Mar 1;56(2):179–83.

26. Patel O, Roy MJ, Kropp A, Hardy JM, Dai W, Lucet IS. Structural basis for small molecule targeting of Doublecortin Like Kinase 1 with DCLK1-IN-1. Commun Biol. 2021 Sep 20;4(1):1105.

27. Cheng L, Yang Z, Guo W, Wu C, Liang S, Tong A, et al. DCLK1 autoinhibition and activation in tumorigenesis. Innovation (Camb). 2022 Jan 25;3(1):100191.

28. Byrne DP, Shrestha S, Galler M, Cao M, Daly LA, Campbell AE, et al. Aurora A regulation by reversible cysteine oxidation reveals evolutionarily conserved redox control of Ser/Thr protein kinase activity. Sci Signal. 2020 Jul 7;13(639):eaax2713.

29. Ferguson FM, Nabet B, Raghavan S, Liu Y, Leggett AL, Kuljanin M, et al. Discovery of a selective inhibitor of doublecortin like kinase 1. Nat Chem Biol. 2020 Jun;16(6):635–43.

30. Neuwald AF. A Bayesian Sampler for Optimization of Protein Domain Hierarchies. Journal of Computational Biology. 2014 Feb 4;21(3):269–86.

31. Rosenberg OS, Deindl S, Sung RJ, Nairn AC, Kuriyan J. Structure of the autoinhibited kinase domain of CaMKII and SAXS analysis of the holoenzyme. Cell. 2005 Dec 2;123(5):849–60.

32. Goldberg J, Nairn AC, Kuriyan J. Structural basis for the autoinhibition of calcium/calmodulin-dependent protein kinase I. Cell. 1996 Mar 22;84(6):875–87.

33. Zheng J, Trafny EA, Knighton DR, Xuong NH, Taylor SS, Ten Eyck LF, et al. 2.2 A refined crystal structure of the catalytic subunit of cAMP-dependent protein kinase complexed with MnATP and a peptide inhibitor. Acta Crystallogr D Biol Crystallogr. 1993 May 1;49(Pt 3):362–5.

34. Romano RA, Kannan N, Kornev AP, Allison CJ, Taylor SS. A chimeric mechanism for polyvalent trans-phosphorylation of PKA by PDK1. Protein Science. 2009;18(7):1486–97.

35. Baffi TR, Newton AC. mTOR Regulation of AGC Kinases: New Twist to an Old Tail. Mol Pharmacol. 2022 Apr 1;101(4):213–8.

36. Taylor SS, Kornev AP. Protein kinases: evolution of dynamic regulatory proteins. Trends Biochem Sci. 2011 Feb;36(2):65–77.

37. Yokokura H, Picciotto MR, Nairn AC, Hidaka H. The regulatory region of calcium/calmodulin-dependent protein kinase I contains closely associated autoinhibitory and calmodulin-binding domains. J Biol Chem. 1995 Oct 6;270(40):23851–9.

38. Yang E, Schulman H. Structural examination of autoregulation of multifunctional calcium/calmodulin-dependent protein kinase II. J Biol Chem. 1999 Sep 10;274(37):26199–208.

39. Eyers PA. TRIBBLES: A Twist in the Pseudokinase Tail. Structure. 2015 Nov 3;23(11):1974–6.

40. Eyers PA, Keeshan K, Kannan N. Tribbles in the 21st Century: The Evolving Roles of Tribbles Pseudokinases in Biology and Disease. Trends Cell Biol. 2017 Apr;27(4):284–98.

41. Foulkes DM, Byrne DP, Yeung W, Shrestha S, Bailey FP, Ferries S, et al. Covalent inhibitors of EGFR family protein kinases induce degradation of human Tribbles 2 (TRIB2) pseudokinase in cancer cells. Sci Signal. 2018 Sep 25;11(549):eaat7951.

42. Harris JA, Fairweather E, Byrne DP, Eyers PA. Analysis of human Tribbles 2 (TRIB2) pseudokinase. Methods Enzymol. 2022;667:79–99.

43. Keeshan K, Bailis W, Dedhia PH, Vega ME, Shestova O, Xu L, et al. Transformation by Tribbles homolog 2 (Trib2) requires both the Trib2 kinase domain and COP1 binding. Blood. 2010 Dec 2;116(23):4948–57.

44. Murphy JM, Nakatani Y, Jamieson SA, Dai W, Lucet IS, Mace PD. Molecular Mechanism of CCAAT-Enhancer Binding Protein Recruitment by the TRIB1 Pseudokinase. Structure. 2015 Nov 3;23(11):2111–21.

45. Burgess HA, Reiner O. Alternative splice variants of doublecortin-like kinase are differentially expressed and have different kinase activities. J Biol Chem. 2002 May 17;277(20):17696–705.

46. Qu D, Weygant N, Yao J, Chandrakesan P, Berry WL, May R, et al. Overexpression of DCLK1-AL Increases Tumor Cell Invasion, Drug Resistance, and KRAS Activation and Can Be Targeted to Inhibit Tumorigenesis in Pancreatic Cancer. J Oncol. 2019;2019:6402925.

47. Ding L, Yang Y, Ge Y, Lu Q, Yan Z, Chen X, et al. Inhibition of DCLK1 with DCLK1-IN-1 Suppresses Renal Cell Carcinoma Invasion and Stemness and Promotes Cytotoxic T-Cell-Mediated Anti-Tumor Immunity. Cancers (Basel). 2021 Nov 16;13(22):5729.

48. Major J, Szende B, Lapis K, Thész Z. Increased SCE inducibility by low doses of methylcholanthrene in lymphocytes obtained from patients with Down’s disease. Mutat Res. 1985 Mar;149(1):51–5.

49. Huang LC, Taujale R, Gravel N, Venkat A, Yeung W, Byrne DP, et al. KinOrtho: a method for mapping human kinase orthologs across the tree of life and illuminating understudied kinases. BMC Bioinformatics. 2021 Sep 18;22(1):446.

50. UniProt Consortium. UniProt: the universal protein knowledgebase in 2021. Nucleic Acids Res. 2021 Jan 8;49(D1):D480–9.

51. Neuwald AF. Rapid detection, classification and accurate alignment of up to a million or more related protein sequences. Bioinformatics. 2009 Aug 1;25(15):1869–75.

52. Nguyen LT, Schmidt HA, von Haeseler A, Minh BQ. IQ-TREE: a fast and effective stochastic algorithm for estimating maximum-likelihood phylogenies. Mol Biol Evol. 2015 Jan;32(1):268–74.

53. Hoang DT, Chernomor O, von Haeseler A, Minh BQ, Vinh LS. UFBoot2: Improving the Ultrafast Bootstrap Approximation. Mol Biol Evol. 2018 Feb 1;35(2):518–22.

54. Le SQ, Gascuel O. An improved general amino acid replacement matrix. Mol Biol Evol. 2008 Jul;25(7):1307–20.

55. Gu X, Fu YX, Li WH. Maximum likelihood estimation of the heterogeneity of substitution rate among nucleotide sites. Mol Biol Evol. 1995 Jul;12(4):546–57.

56. Kalyaanamoorthy S, Minh BQ, Wong TKF, von Haeseler A, Jermiin LS. ModelFinder: fast model selection for accurate phylogenetic estimates. Nat Methods. 2017 Jun;14(6):587–9.

57. Huerta-Cepas J, Serra F, Bork P. ETE 3: Reconstruction, Analysis, and Visualization of Phylogenomic Data. Mol Biol Evol. 2016 Jun;33(6):1635–8.

58. Katoh K, Misawa K, Kuma K ichi, Miyata T. MAFFT: a novel method for rapid multiple sequence alignment based on fast Fourier transform. Nucleic Acids Res. 2002 Jul 15;30(14):3059–66.

59. Mandell DJ, Coutsias EA, Kortemme T. Sub-angstrom accuracy in protein loop reconstruction by robotics-inspired conformational sampling. Nat Methods. 2009 Aug;6(8):551–2.

60. Tyka MD, Keedy DA, André I, Dimaio F, Song Y, Richardson DC, et al. Alternate states of proteins revealed by detailed energy landscape mapping. J Mol Biol. 2011 Jan 14;405(2):607–18.

61. Kabsch W, Sander C. Dictionary of protein secondary structure: pattern recognition of hydrogen-bonded and geometrical features. Biopolymers. 1983 Dec;22(12):2577–637.

62. Michaud-Agrawal N, Denning EJ, Woolf TB, Beckstein O. MDAnalysis: a toolkit for the analysis of molecular dynamics simulations. J Comput Chem. 2011 Jul 30;32(10):2319–27.

63. Jorgensen WL, Chandrasekhar J, Madura JD, Impey RW, Klein ML. Comparison of simple potential functions for simulating liquid water. J Chem Phys. 1983 Jul 15;79(2):926–35.

64. Pronk S, Páll S, Schulz R, Larsson P, Bjelkmar P, Apostolov R, et al. GROMACS 4.5: a high-throughput and highly parallel open source molecular simulation toolkit. Bioinformatics. 2013 Apr 1;29(7):845–54.

65. Huang J, MacKerell AD. CHARMM36 all-atom additive protein force field: validation based on comparison to NMR data. J Comput Chem. 2013 Sep 30;34(25):2135–45.

66. Humphrey W, Dalke A, Schulten K. VMD: visual molecular dynamics. J Mol Graph. 1996 Feb;14(1):33–8, 27–8.

67. Park H, Bradley P, Greisen P, Liu Y, Mulligan VK, Kim DE, et al. Simultaneous Optimization of Biomolecular Energy Functions on Features from Small Molecules and Macromolecules. J Chem Theory Comput. 2016 Dec 13;12(12):6201–12.

68. Keller O, Odronitz F, Stanke M, Kollmar M, Waack S. Scipio: using protein sequences to determine the precise exon/intron structures of genes and their orthologs in closely related species. BMC Bioinformatics. 2008 Jun 13;9:278.

69. Cunningham F, Allen JE, Allen J, Alvarez-Jarreta J, Amode MR, Armean IM, et al. Ensembl 2022. Nucleic Acids Res. 2022 Jan 7;50(D1):D988–95.

70. Ferries S, Perkins S, Brownridge PJ, Campbell A, Eyers PA, Jones AR, et al. Evaluation of Parameters for Confident Phosphorylation Site Localization Using an Orbitrap Fusion Tribrid Mass Spectrometer. J Proteome Res. 2017 Sep 1;16(9):3448–59.

71. Daly LA, Brownridge PJ, Batie M, Rocha S, Sée V, Eyers CE. Oxygen-dependent changes in binding partners and post-translational modifications regulate the abundance and activity of HIF-1α/2α. Sci Signal. 2021 Jul 20;14(692):eabf6685.

72. Silva JC, Gorenstein MV, Li GZ, Vissers JPC, Geromanos SJ. Absolute quantification of proteins by LCMSE: a virtue of parallel MS acquisition. Mol Cell Proteomics. 2006 Jan;5(1):144–56.

73. Byrne DP, Clarke CJ, Brownridge PJ, Kalyuzhnyy A, Perkins S, Campbell A, et al. Use of the Polo-like kinase 4 (PLK4) inhibitor centrinone to investigate intracellular signalling networks using SILAC-based phosphoproteomics. Biochem J. 2020 Jul 17;477(13):2451–75.

74. Byrne DP, Vonderach M, Ferries S, Brownridge PJ, Eyers CE, Eyers PA. cAMP-dependent protein kinase (PKA) complexes probed by complementary differential scanning fluorimetry and ion mobility-mass spectrometry. Biochem J. 2016 Oct 1;473(19):3159–75.

75. Omar MH, Kihiu M, Byrne DP, Lee KS, Lakey TM, Butcher E, et al. Classification of Cushing’s syndrome PKAc mutants based upon their ability to bind PKI. Biochem J. 2023 Jun 28;480(12):875– 90.

76. McSkimming DI, Dastgheib S, Baffi TR, Byrne DP, Ferries S, Scott ST, et al. KinView: a visual comparative sequence analysis tool for integrated kinome research. Mol Biosyst. 2016 Nov 15;12(12):3651–65.

